# Coral accurately bridges paired-end RNA-seq reads alignment

**DOI:** 10.1101/2020.03.03.975821

**Authors:** Qian Shi, Mingfu Shao

## Abstract

**Motivation:** The established high-throughput RNA-seq technologies usually produce paired-end reads. A challenging problem is therefore to computationally infer the alignment of entire fragments given the alignment of the two mate ends. Solving this problem essentially provide longer RNA-seq reads, and hence benefits downstream RNA-seq analysis.

**Results:** We introduce Coral, a new tool that can accurately bridge paired-end RNA-seq reads. The core of Coral is a novel optimization formulation that can capture the most reliable bridging path while also filter out false paths. An efficient dynamic programming algorithm is designed to calculate the top *N* optimum. Coral implements a consensus approach to select the best solution among the *N* candidates by taking into account the distribution of fragment length. Coral is modular, can be easily incorporated into existing RNA-seq analysis pipeline. We show that Coral can improve transcript assembly by a large margin: on average over 2377 RNA-seq samples from GTEx, the improvement (measured with adjusted precision) is 7.5% and 11.2% when Coral is incorporated with StringTie and Scallop, respectively.

**Availability:** Coral is open-source, freely available at GitHub (https://github.com/Shao-Group/coral) and Bioconda. Scripts, datasets and documentations that can reproduce all experimental results in this paper are available at https://github.com/Shao-Group/coraltest.

## 1 Introduction

The established high-throughput RNA sequencing technologies (RNA-seq) enables global and accurate measurement of isoform-level gene activities. The second generation RNA-seq (i.e., short-reads RNA-seq) usually produces *paired-end* reads, which report the sequences of the two ends of a fragment, while ignore the middle portion of the fragment (if the two ends cannot cover the entire fragment). The fact that the two ends are from the same fragment and that the length of fragments follows a certain distribution (through fragment size selection) provide valuable long-range information in determining complicated splicing variants. Such paired-end information has been widely used in various RNA-seq analysis tasks to improve accuracy, including splicing-aware alignment (e.g., STAR [1], HISAT [2], SpliceMap [3]), expression quantification (e.g., Salmon [4], kallisto [5], RSEM [6]), assembly (e.g., StringTie [7], TransComb [8], Scallop [9]), fusion detection (e.g., FuSeq [10], STAR-Fusion [11], SQUID [12]), and splicing quantification (e.g., DARTS [13], leafCutter [14]), among many other tools and software.

We explore the problem of computationally inferring the *entire* fragments given their sequenced two ends. We do this in a reference-based setting: given the alignment of the two sequenced ends to a reference genome, to infer the alignment of entire fragments. Such computational inference is possible, as we expect that most of the splicing junctions can be detected by spliced reads (a reference annotation can supplement missing splicing junctions to some extent), and the alignment of entire fragments can be obtained by traversing the splicing junctions to bridge their two ends. More specifically, the splicing junctions can be organized with a *splice graph*, in which each vertex corresponds to a (partial) exon and each edge corresponds to a splicing junction (see details in Methods and Figure 1). The alignment of the two mate ends *e*_1_ and *e*_2_ of a fragment can then be represented as two paths *p*_1_ and *p*_2_ in the splice graph, and the alignment of the entire fragment can be inferred by finding an appropriate path *p* that connects *p*_1_ and *p*_2_ in the splice graph (path connecting *p*_1_, *p*, and *p*_2_ gives the alignment of the entire fragment).

**Figure 1:**
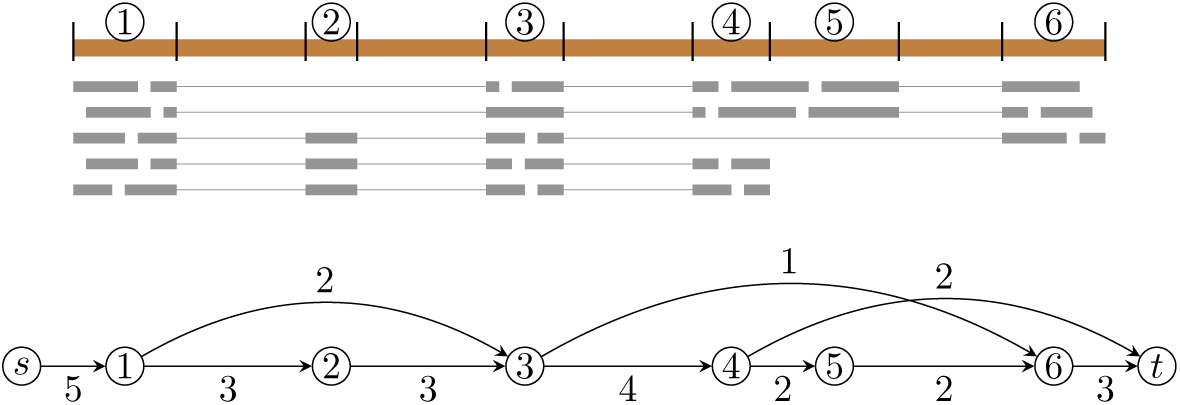
Example of the reads alignment in a gene locus (upper part). Reference genome are portioned into partial exons (numbered from 1 to 6) using the splicing coordinates. Lower part shows the corresponding splice graph; the abundance of each edge is given next.

Several challenges exist in above computational inference. First, because of the complicated mechanism of alternative splicing and dynamic nature of transcription and mRNA degradation, many different types of splicing junctions will be captured by RNA-seq reads. According to our experiments, in many gene loci, thousands of different splicing junctions can be observed (a majority of them are with low abundance). The number of possible bridging paths are exponential in the number of junctions, which results in a huge search space when determining the optimal solution. Second, sequencing errors and alignment errors produce false junctions, which therefore introduces false bridging paths and further enlarges the search space. Third, it remains open what is a good characterization of the true bridging paths; in other words what the objective function should be when this task is formulating as an optimization problem. Fourth, the distribution of fragment length provides information in deciding false bridging paths, but such information is noisy and weak, and how to make use of it remain challenging.

Although many RNA-seq analysis tools use paired-end information in some way, limited efforts have been made to directly and explicitly infer the alignment of entire fragments. Existing method for this purpose includes MapPER [15]. MapPER implements a probabilistic framework: starting from splicing junctions of all end reads, MapPER constructs potential splicing paths connecting paired-end reads; an expectation maximization method assigns likelihood values to all splice junctions and assigns the most probable alignment for each fragment. MapPER has been shown improving junction detection.

In this paper, we present a new method, called *Coral*, for bridging paired-end RNA-seq reads. In Section 2, we propose a novel formulation for this task and devise an efficient algorithm. We also design a new method using voting to make use of the length distribution to obtain more robust inference. In Section 3, we use extensive experimental studies to show that Coral can significantly improve transcript assembly. Conclusion and future directions are described in Sections 4 and 5.

## 2 Methods

We formulate the task of bridging paired-end reads as a new optimization problem, and designed an efficient algorithm. In Section 2.1, we describe the formulation and the algorithm. In Section 2.2, we describe its extension when the reference annotation is provided. In Section 2.3, we describe a new method that can correct the errors in the alignments based on the algorithm given in Section 2.1. Throughout this section, we use “read” to refer to an individual sequencing end, and use “fragment” to refer to a pair of mate ends (i.e., paired-end reads).

### 2.1 Formulation and Algorithm

The input for Coral is standard RNA-seq alignment in sam/bam format. Coral first groups fragments into gene loci: reads that overlap on their alignment coordinates will be assigned to the same gene locus, and Coral makes sure that the two ends from the same fragment (which will have the same ID in the sam/bam file) will be assigned to the same gene locus. Each gene locus will be treated as independent instances and solved independently with the algorithm given below.

In each gene locus, junctions will be extracted from spliced reads. The collected set of splicing coordinates will be used to partition the reference genome into continuous segments, called *partial exons*. We use the well-known *splice graph* to represent the partial exons and junctions in a single gene locus (see Figure 1): each partial exon corresponds to a vertex, and each junction corresponds to a direct edge that connects its two corresponding exons. Two additional vertices, source *s* and sink *t*, are added and connected to possible starting and ending partial exons. The *abundance* of an edge is calculated as the number of reads that contains the corresponding junction.

With the splice graph, each read will be then represented as the list of the vertices to which it is aligned, and each fragment will be then represented as a pair of lists corresponding to its two ends. We then cluster all fragments into *equivalent classes*: fragments that are represented as exactly the same pair of lists from an equivalent class. An equivalent class is also represented as a pair of lists of vertices of the splice graph.

Let *G* = (*V, E*) be the corresponding splice graph of a gene locus. Let *A* = ((*a*_1_, *a*_2_, …, *a*_*m*_), (*b*_1_, *b*_2_, …, *b*_*m*′_)) be an equivalent class. The problem of bridging a fragment *f* in *A* is therefore to find a path from *a*_*m*_ to *b*_1_ in the splice graph *G*. Such path, called *bridging path*, infers the alignment of the unsequenced portion of fragment *f*. We assume that, all fragments in *A* have identical bridging path. This is because fragments in an equivalent class are similar, as their two ends are aligned to the same list of vertices. Algorithmically, this assumption allows us to reduce computational efforts (as all fragments can be bridged in a single run, rather than bridging them individually) while also enables a robust voting scheme to select the best bridging path from a set of candidates by integrating distribution of fragment length (see below).

Our core algorithm to bridge all fragments in an equivalent class *A* = ((*a*_1_, *a*_2_, …, *a*_*m*_), (*b*_1_, *b*_2_, …, *b*_*m′*_)) consists of two steps, *nominating* and *voting*. The nominating step computes a set of candidate bridging paths from *a*_*m*_ to *b*_1_ in *G*, and the voting step selects one through a consensus approach with the fragment length distribution.

We formulate the nominating step as a new optimization problem and then design an efficient algorithm. We start from defining a full ordering for all possible bridging paths. Let *p*_1_ and *p*_2_ be two arbitrary paths from *a*_*m*_ to *b*_1_ in *G*. Let 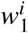 (resp. 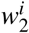) be the *i*th smallest edge abundance among all edges in path *p*_1_ (resp. *p*_2_). We say *p*_1_ is *more reliable* than *p*_2_, if there exists an integer *k* such that 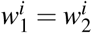, for all 1 ≤ *i < k*, and 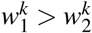. (More formally, each bridging path is represented as the sorted list of its edge abundances in ascending order; then bridging paths are sorted in lexicographical order.) Our formulation for the nominating step is to find *N* most reliable paths from *a*_*m*_ to *b*_1_ in *G* (i.e., the first *N* bridging paths in the lexicographical order), where *N* is parameter of Coral (default value is 10).

The intuition of above formulation is to find a set of bridging paths such that the abundances of their *bottle-neck* edges (i.e., edges with smallest abundances) are as large as possible (if the smallest abundances are the same, we select ones whose second smallest is maximized, and so on). This formulation has two advantages. First, by maximizing bottleneck abundances, the bridging paths are supported strongest, and hence are more likely to be true bridging path. Second, through this formulation, paths with false splicing junctions, which are usually due to alignment errors and have low abundances, can be automatically excluded, and therefore false bridging paths can be efficiently avoided.

This formulation satisfies the *optimal substructure* property: if 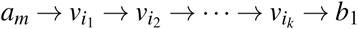 is the most reliable path from *a*_*m*_ to *b*_1_, then 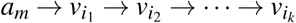 is the most reliable path from *a*_*m*_ to 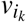. Hence, a standard dynamic programming can be designed to find the optimal solution. We use an algorithm that computes *N* most reliable paths for all pairs of vertices in *G*. (They will be stored and be fetched as needed when processing each equivalent class.) Specifically, let (*v*_1_, *v*_2_, …, *v*_*n*_) be a topological sorting of *V* (this is possible because splice graph is a directed acyclic graph). Given a particular *v*_*i*_, we can use a single run to find *N* most reliable paths from *v*_*i*_ to *v* _*j*_ for every *j* > *i*. To compute these *N* paths for *v* _*j*_, we examine all vertices *v*_*k*_ that directly connects to *v* _*j*_, and compare all paths stored in these vertices (each *v*_*k*_ already stores *N* most reliable paths from *v*_*i*_ to *v*_*k*_ at this time point). The best *N* of them will be kept and after concatenating *v* _*j*_ they become the *N* most reliable paths from *v*_*i*_ to *v* _*j*_. We run this subroutine for all *v*_*i*_, 1 ≤ *i* ≤ *n*, which gives *N* most reliable paths for all pairs of vertices in *G*. The overall running time of this algorithm is *O*(*N* · |*V*|^2^ · |*E*|). To speed up, instead of maintaining the full list of the edge abundances for each path, whose length is *O*(|*V*|), in Coral we only store the smallest *M* edge abundances (*M* is a parameter with default value of 5). This gives an improved running time of *O*(*N* · *M* · |*V* | · |*E*|). Although optimality may not be guaranteed, experimental studies show that this heuristic rarely affects the overall accuracy.

We now describe the voting step. Let *A* be an equivalent class defined above. The above nominating algorithm gives us *N* candidate bridging paths {*p*_1_, *p*_2_, …, *p*_*N*_} from *a*_*m*_ to *b*_1_. Assume that these *N* paths are sorted (i.e., *p*_*i*_ is more reliable than *p*_*i*+1_). We now examine each fragment *f* in *A*. For each candidate path *p*_*i*_, 1 ≤ *i* ≤ *N*, we calculate the *actual sequence length* of fragment *f*, denoted as *f*_*i*_, assuming that *f* is bridged with *p*_*i*_. This can be done because once the bridging path of *f* is given, the alignment of the *entire* sequence of fragment *f* is fixed and hence the length of its entire sequence can be calculated. If *f*_*i*_ is within a *reasonable* range that precomputed in the very beginning of Coral (100,000 paired-end reads with unique bridging path will be sampled to compute the empirical distribution of fragment length, and the value from percentile 2 to percentile 98 will be used as the reasonable range), then fragment *f* votes path *p*_*i*_; otherwise, we try next best path (i.e., *p*_*i*+1_) and check whether *f*_*i*+1_ is within the reasonable range. In other words, each fragment votes the best candidate that leads to a fragment length within the reasonable range. Candidate with the largest number of votes will be accepted as the bridging path for the equivalent class, and all fragments in it will be bridged using this path. The alignment of the entire fragment will be determined following this bridging path and written into the resulting sam/bam file. If none of the candidate paths receives any vote, then this equivalent class fails bridging and the original alignments of the individual reads will be reported without changing.

### 2.2 Incorporating Reference Annotation

Coral supports using the reference annotation to improve accuracy, as true junctions might be missed due to low coverage, sequencing and alignment errors. If the reference annotation, i.e., a set of known transcripts, is provided, Coral will first find candidate bridging paths solely based on the annotations (instead of running the nominating step described in Section 2.1). Specifically, for each equivalent class *A* = (*a*_1_, *a*_2_, …, *a*_*m*_), (*b*_1_, *b*_2_, …, *b*_*m*′_)), Coral locates the set of known transcripts *T* spans both (*a*_1_, *a*_2_, …, *a*_*m*_) and (*b*_1_, *b*_2_, …, *b*_*m*′_), i.e., each transcript in *T* can bridge all fragments in *A*. To speed up this process, Coral will first create an index for all known transcripts so that such *T* can be quickly calculated. After having the candidate bridging paths, the same voting step will be used to select the final bridging path. If *T* is empty or none of the transcripts in *T* gets any vote, the algorithm described in Section 2.1 will be then applied.

### 2.3 Correcting Alignment Errors

We propose a novel method to correct alignment errors while bridging (Figure 2). Reads that span a junction but flank one side shortly are prone to alignment errors (left red read in Figure 2). Information of paired-end can be used to detect and correct such errors. Specifically, Let *A* = ((*a*_1_, *a*_2_, …, *a*_*m*_), (*b*_1_, *b*_2_, …, *b*_*m*′_)) be an equivalent class that is failed bridging. If the length of the aligned portion on *a*_*m*_ is small (a tunable parameter with default value of 10 basepairs), Coral will then try to bridge *A*_1_ = (*a*_1_, *a*_2_, …, *a*_*m*-1_), (*b*_1_, *b*_2_, …, *b*_*m*′_)). Symmetrically, Coral will also examine the length of the aligned portion on *b*_1_ and examine whether *A*_2_ = ((*a*_1_, *a*_2_, …, *a*_*m*_), (*b*_2_, *b*_3_, …, *b*_*m*′_)) or *A*_3_ = ((*a*_1_, *a*_2_, …, *a*_*m*-1_), (*b*_2_, *b*_3_, …, *b*_*m*′_)) can be successfully bridged. If so, the corrected alignment will be reported.

**Figure 2:**
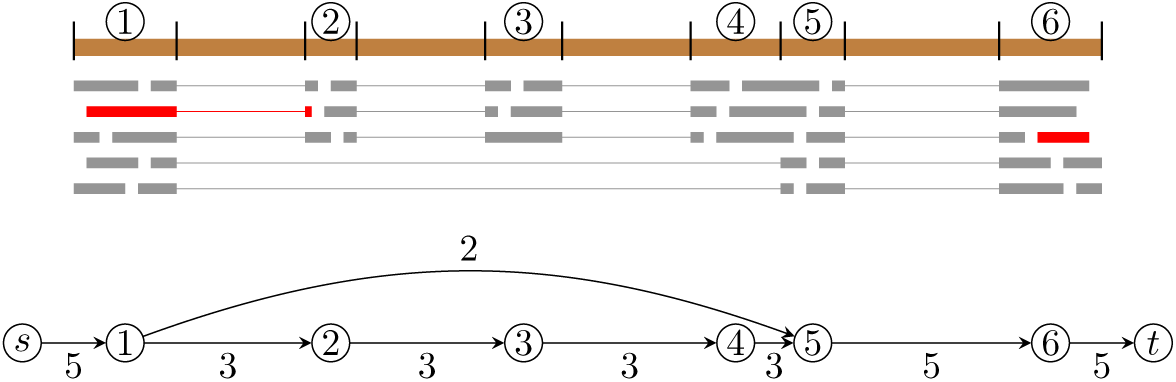
Example of correcting alignments while bridging. One fragment (in red) is in equivalent class ((1,2), (6)). The only bridging path is (2,3,4,5,6), which is most likely beyond the reasonable range of fragment length. As the aligned portion on segment 2 of the first-end is short, Coral will shorten (1,2) as (1) and then try to bridge it in equivalent class ((1), (6)), for which there is an additional candidate (1,5,6), which gives this fragment a sequence length within the reasonable range. Therefore, Coral will report (1,5,6) as the resulting alignment for this paired-end read, suggesting that the alignment of its first-end is wrong (its short flanking region should be aligned to segment 5).

## 3 Results

Coral infers the alignment of entire fragments, essentially providing longer aligned reads that can improve downstream analysis. Such information is particularly useful in resolving complicated splicing variants of the expressed transcripts. Below, we extensively evaluate the effectiveness of Coral in improving transcript assembly. The scripts and detailed description on how to reproduce the experimental results and figures in this paper are available at https://github.com/Shao-Group/coraltest.

We follow the workflow illustrated in Figure 3 to evaluate the effectiveness of Coral in improving transcript assembly. This workflow is configured by 3 factors: the RNA-seq aligner (we experiment two widely-used aligners STAR [1] and HISAT [2]), the transcript assembler (we test two recent and leading assemblers StringTie [7] and Scallop [9]), and whether a reference transcriptome is provided to Coral (see Section 2.2).

**Figure 3:**
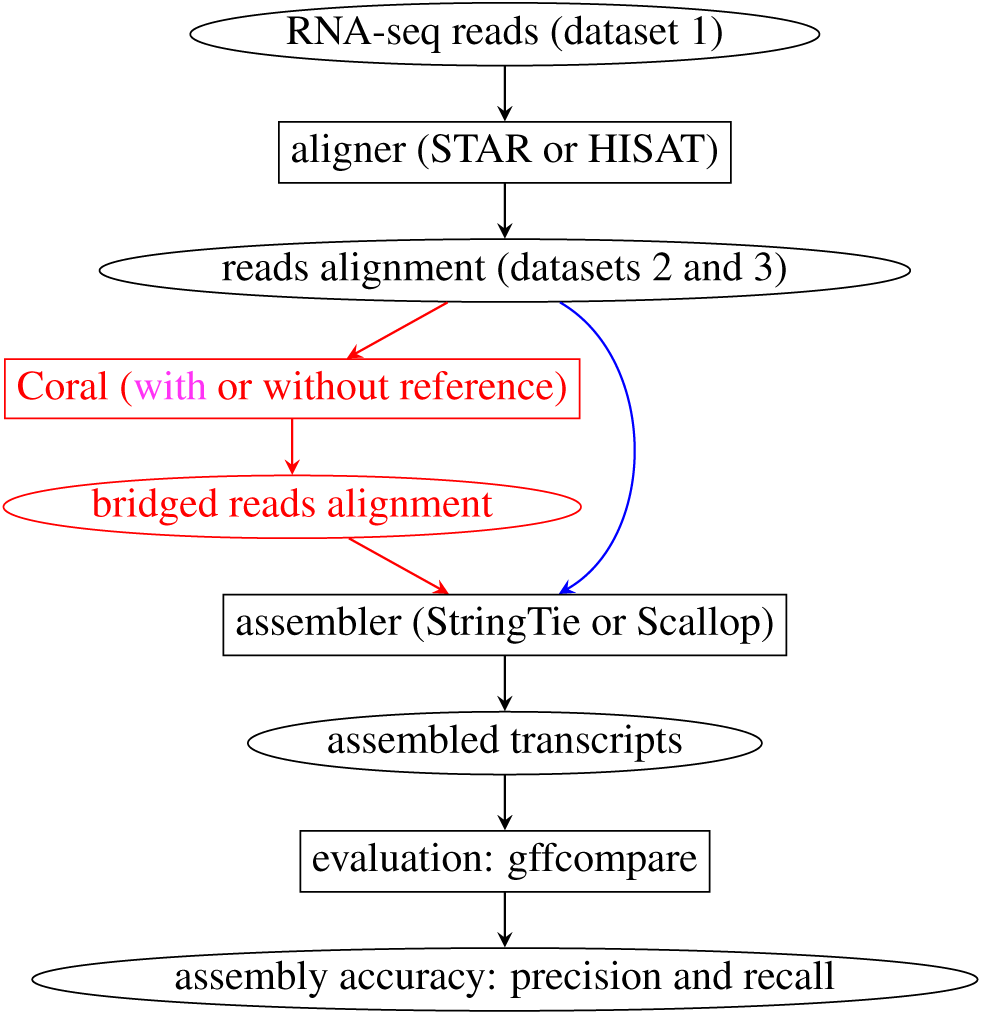
Workflow of evaluating the effectiveness of Coral in improving transcript assembly. The three configurations without and with using Coral (whether reference is further provided) are marked blue and red, and rose.

For each workflow configuration (choice of aligner and assembler), we compare the assembly accuracies of the three configurations: without Coral, with Coral and Coral is not provided with reference annotation and with Coral and Coral is provided with reference annotation (see blue and red parts of Figure 3). For each configuration, the assembler predicts a set of transcripts. We define a predicted transcript is *known* if its intron-chain coordinates exactly match those of an existing transcript in the reference transcriptome. We reports two measures, the number of known transcripts (proportional to sensitivity), and precision (defined as the ratio between the number of known transcripts and the total number of predicted transcripts). We use a third-party tool, gffcompare, to compute these two measures. We use these measures for assembly to evaluate the effectiveness of Coral in improving transcript assembly.

On some samples one configuration obtains higher number of known transcripts but lower precision, or vise versa. To still compare the three configurations on such samples, we first balance them in one measure and then compare the other. Specifically, let *k*_1_, *k*_2_, and *k*_3_ be the number of known transcripts obtained by the three configurations *X*_1_, *X*_2_, and *X*_3_, and *p*_1_, *p*_2_ and *p*_3_ be their corresponding precisions. Assume that *k*_1_ > *k*_2_ > *k*_3_. We sort the assembled transcripts obtained by *X*_1_ based on the expression abundance, and gradually filter out assembled transcripts with lowest abundance. In this way, the known transcripts of *X*_1_ will decrease while its precision will likely increase (as lowly-expressed ones are more likely to be false positive transcripts). We calculate the corresponding new measures of *X*_1_, denoted as 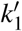 and 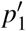, as filtering goes, and stop when 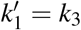. We then report 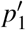 at this time as the *adjusted precision* of *X*_1_ (now the number of known transcripts of *X*_1_ becomes 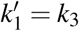, the same as *X*_3_). We do exactly the same balancing for *X*_2_ to get its adjusted precision of 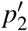. Note that the relationship between 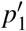, and 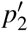 and *p*_3_ can therefore reflect the overall performance of the three configurations, as their number of known transcripts have been adjusted identical. Similarly, we also compute the *adjusted known* transcripts, through gradually filtering out the transcripts with lowest abundance in the two configurations with lower original precision, until their (adjusted) precision matches the configuration with highest original precision. This way of comparing assembly accuracy has also been used in the Scallop paper [9].

We use 3 datasets to evaluate Coral following above workflow and evaluation criterion. The first two datasets have been used in the Scallop paper [9]: the first dataset contains 10 strand-specific RNA-seq samples from ENCODE, and we test them using different aligners (STAR and HISAT); the second dataset contains 50 strand-specific RNA-seq samples (already aligned) from ENCODE. The third dataset contains 2377 non-strand-specific RNA-seq samples (already aligned) from GTEx.

The original assembly accuracy of the three configurations (i.e., without using Coral, using Coral and run without reference annotation, and using Coral and run with reference annotation) averaged over the 10 samples in dataset 1 is illustrated in Figure 4. Coral improves both recall (proportional to the number of known transcripts) and precision on all combinations of aligners and assemblers used except in one case: for the combination of HISAT and Scallop, the precision gets slightly worse (32.2% vs. 31.4%); notice that for this combination, the recall gets better, and when one of them is balanced, both the adjusted precision and the adjusted recall get improved (Figure 5), suggesting that Coral improves the average transcript assembly for this combination (and therefore for all combinations).

**Figure 4:**
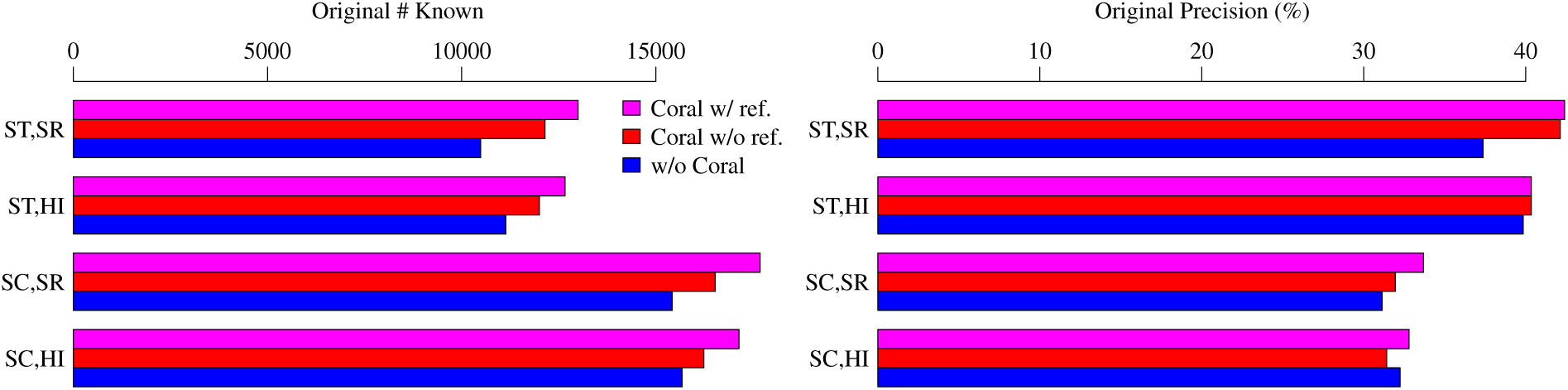
Comparison of the average original assembly accuracy obtained with the 3 configurations over the 10 samples in dataset 1. Abbreviations: ST = StringTie; SC = Scallop; SR = STAR; HI = HISAT.

**Figure 5:**
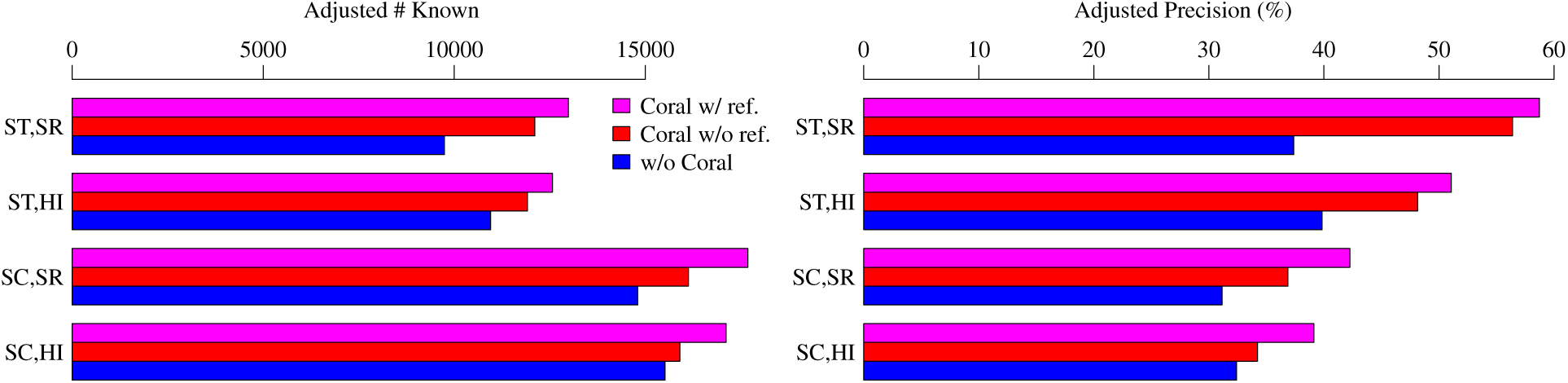
Comparison of the average adjusted assembly accuracy obtained with the 3 configurations over the 10 samples in dataset 1. ST = StringTie; SC = Scallop; SR = STAR; HI = HISAT.

The improvement by using Coral, evaluated with adjusted measures on dataset 1, is illustrated in Figure 5. Coral, when run without reference annotation, can improve transcript assembly on all combinations of aligners and assemblers: the improvement in the average adjusted number of known transcripts, computed as the *ratio* between the two configurations, ranges from 2.5% (HISAT + Scallop) to 24.3% (STAR + StringTie), and in the average adjusted precision the improvement (computed as the *difference* between the two configurations) ranges from 1.8% (HISAT + Scallop) to 19.0% (STAR + StringTie). We note that, when Coral is run without reference annotation, the only input of Coral is the given reads alignment (see Section 2.1), and hence such improvement is purely due to the bridging algorithm within Coral. When Coral is run with the reference annotation, the improvement is more pronounced: in the average adjusted number of known transcripts the improvement ranges from 10.3% to 33.3%, and in the average adjusted precision it ranges from 6.7% to 21.4%. This shows that the additional information of annotations together with the bridging algorithm that utilize it (described in Section 2.2) can further improve bridging paired-end reads.

The performance on individual samples in dataset 1 is shown in Supplementary Figures 1–8. When reference annotation is not provided to Coral, Coral improves on 38 out of 40 cases considering the 4 combinations of aligners and assemblers: the improvement in adjusted precision (i.e., absolute increase) ranges from - 2.6% to 24.0%, and the improvement in the number of adjusted known transcripts (i.e., ratio) ranges from -2.3% to 57.0%. When reference annotation is provided to Coral, Coral improves on 39 out of 40 cases: the improvement in adjusted precision ranges from -0.49% to 28.2%, and the improvement in the number of adjusted known transcripts ranges from -0.59% to 76.2%.

The experimental results on the 50 samples in dataset 2 are given in Figure 6 (average original accuracy), Figure 7 (average adjusted accuracy), and Supplementary Figures 9–16 (accuracy on individual samples). Again assembly accuracy gets improvement when Coral is used in all combinations. For examples, when Coral is not provided with reference annotation, the average adjusted number of known transcripts gets increased 7.1% and 6.8% for assembler StringTie and Scallop respectively (the numbers are 9.7% and 10.6% when reference is used in Coral). Over the 100 individual cases (50 samples with 2 assemblers), 98 of them gets improved when Coral (without reference) is used, ranging from -0.8% to 25.7% in the adjusted number of known transcripts (the range is from 0.5% to 40.3% when reference is used in Coral, i.e., all cases gets improved).

**Figure 6:**
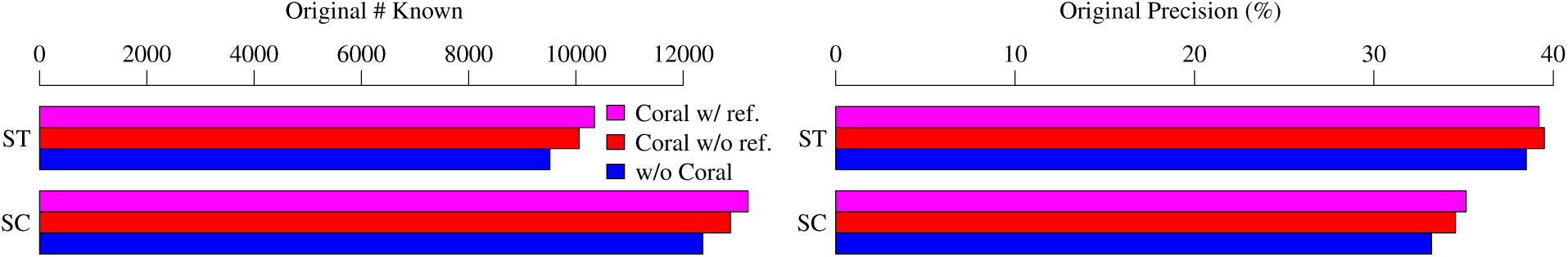
Comparison of the average original assembly accuracy obtained with the 3 configurations over the 50 samples in dataset 2. ST = StringTie; SC = Scallop.

**Figure 7:**
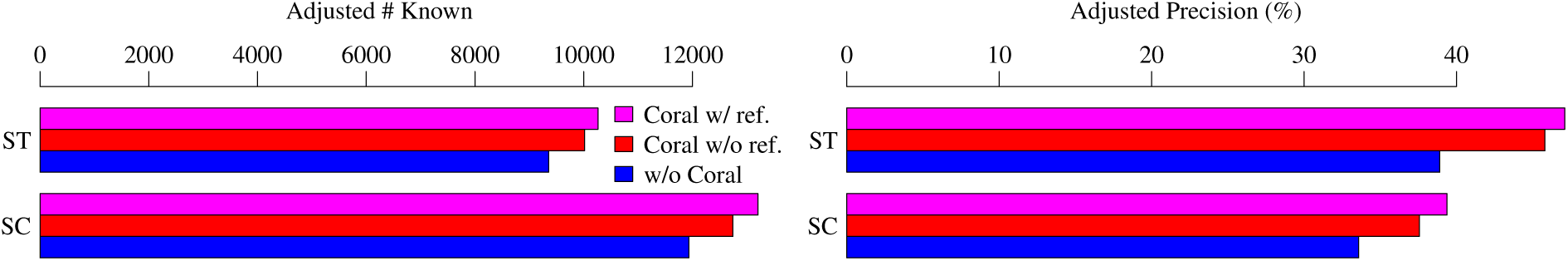
Comparison of the average adjusted assembly accuracy obtained with the 3 configurations over the 50 samples in dataset 2. ST = StringTie; SC = Scallop.

The experimental results on the 2377 RNA-seq samples in dataset 3 are given in Figure 8 (average original accuracy), Figure 9 (average adjusted accuracy), and Supplementary Figures 17–24 (distribution of assembly accuracy). Similar improvement has been obtained. When Coral is not provided with reference annotation, the average adjusted number of known transcripts gets increased 4.9% and 6.1% for assembler StringTie and Scallop respectively (the numbers are 7.5% and 11.2% when reference is used in Coral). Over the 4754 individual cases, *all* of them get improved when Coral (without reference) is used, ranging from 0.8% to 21.7% in the adjusted number of known transcripts (the range is from 1.3% to 34.0% when reference is used in Coral).

**Figure 8:**
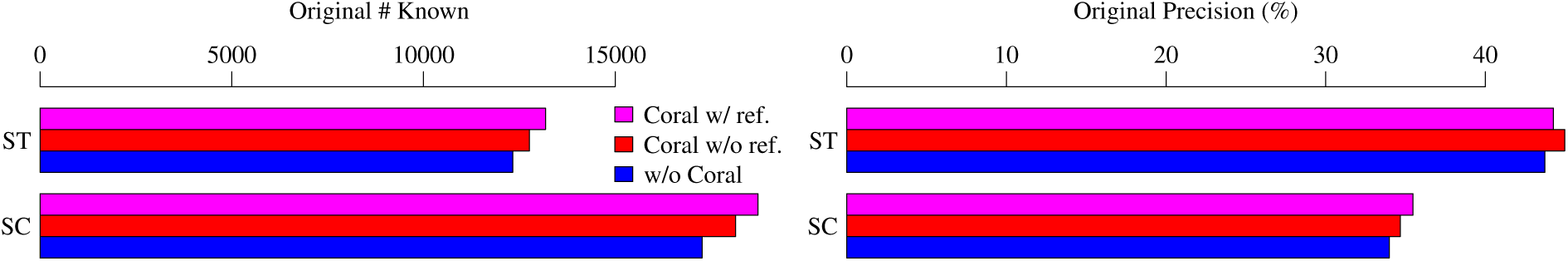
Comparison of the average original assembly accuracy obtained with the 3 configurations over the 2377 samples in dataset 3. ST = StringTie; SC = Scallop.

**Figure 9:**
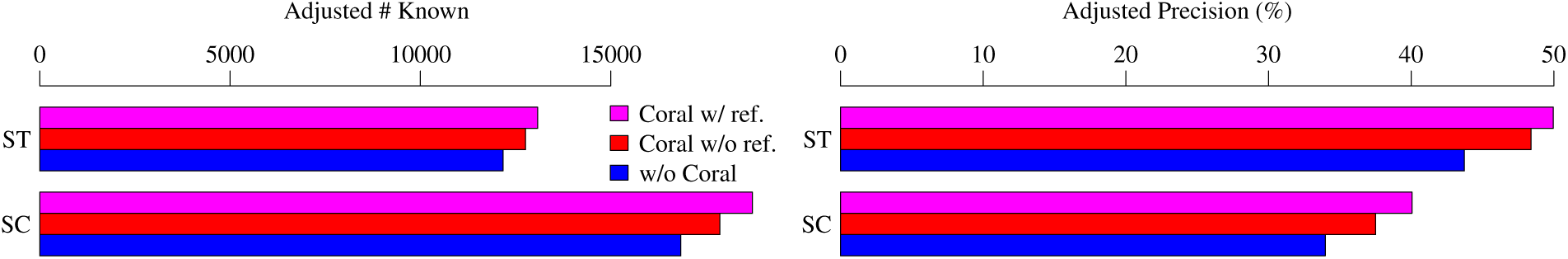
Comparison of the average adjusted assembly accuracy obtained with the 3 configurations over the 2377 samples in dataset 3. ST = StringTie; SC = Scallop.

Coral is fast: the average running time of Coral over the 2377 GTEx samples is 19 minutes (single thread). The distribution is given in Figure 10.

**Figure 10:**
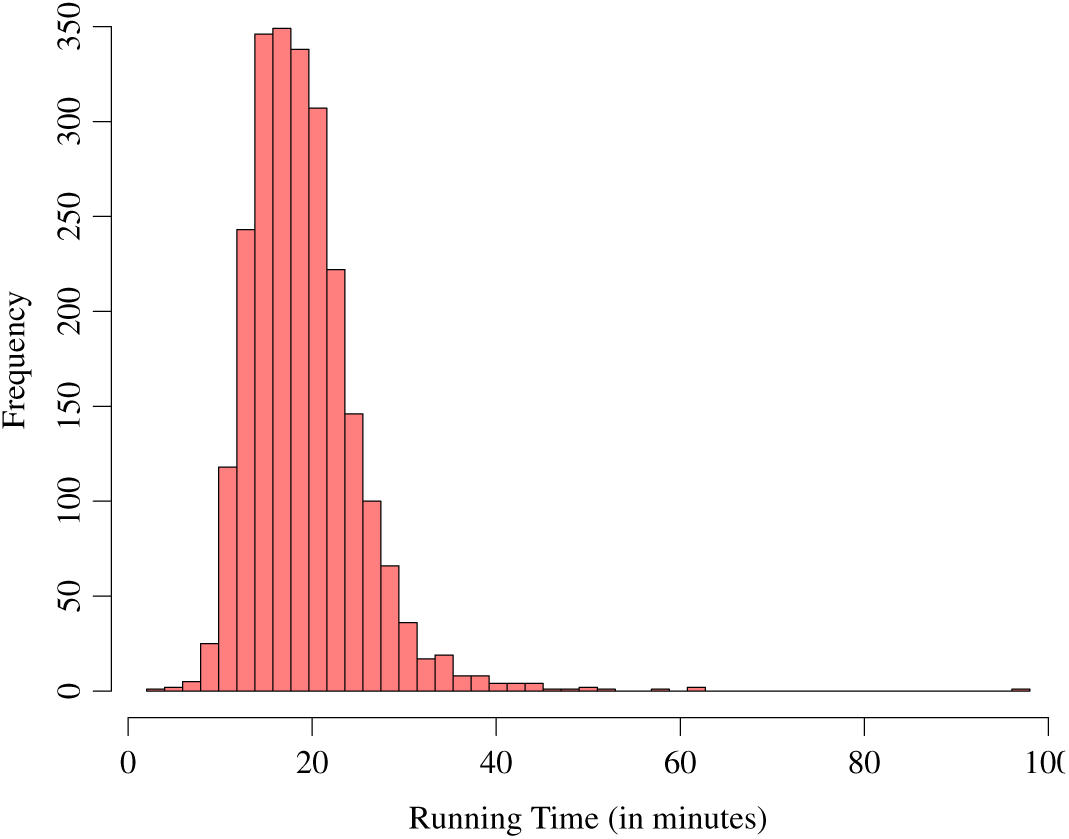
Distribution of the running time of Coral on the 2377 GTEx RNA-seq samples.

## 4 Conclusion

We present Coral, a new tool that can accurately bridge paired-end RNA-seq reads. Our main algorithmic contribution is a novel optimization formulation (i.e., to find bridging paths with maximized bottleneck abundances) that leads to robust inference. A new consensus approach is introduced to further select from the candidates by using the length distribution information. Combined, Coral can efficiently infer the alignment of entire fragments.

Coral is modular, with both input (alignment of paired-end reads) and output (alignment of entire fragments) being standard sam/bam format. It can be very easily incorporated into existing RNA-seq analysis pipeline (i.e., adding one line `./coral -i input.bam -o output.bam`). Coral is available at GitHub and can also be easily installed with Bioconda. We devote to reproducibility: all the experimental results in this paper (and supplementary material) can be reproduced with the scripts available at GitHub.

## 5 Discussion

We showed the effectiveness of Coral in improving transcript assembly. We expect that together with other RNA-seq analysis tools, Coral will be able to improve other downstream RNA-seq analysis, for example, isoform-level quantification and splicing quantification. More specifically, with the inferred alignment of entire fragments, fragments will be less ambiguously aligned to multiple transcripts, and hence improve isoform quantification. Also, with the inferred alignment of fragments, we know the missing junctions in the unsequenced portion of the fragment, which can give more accurate estimation of junction abundance (and hence improve splicing quantification).

