## Supplementary Figures for "Coral accurately bridges paired-end RNA-seq reads alignment"

### Supplementary Materials

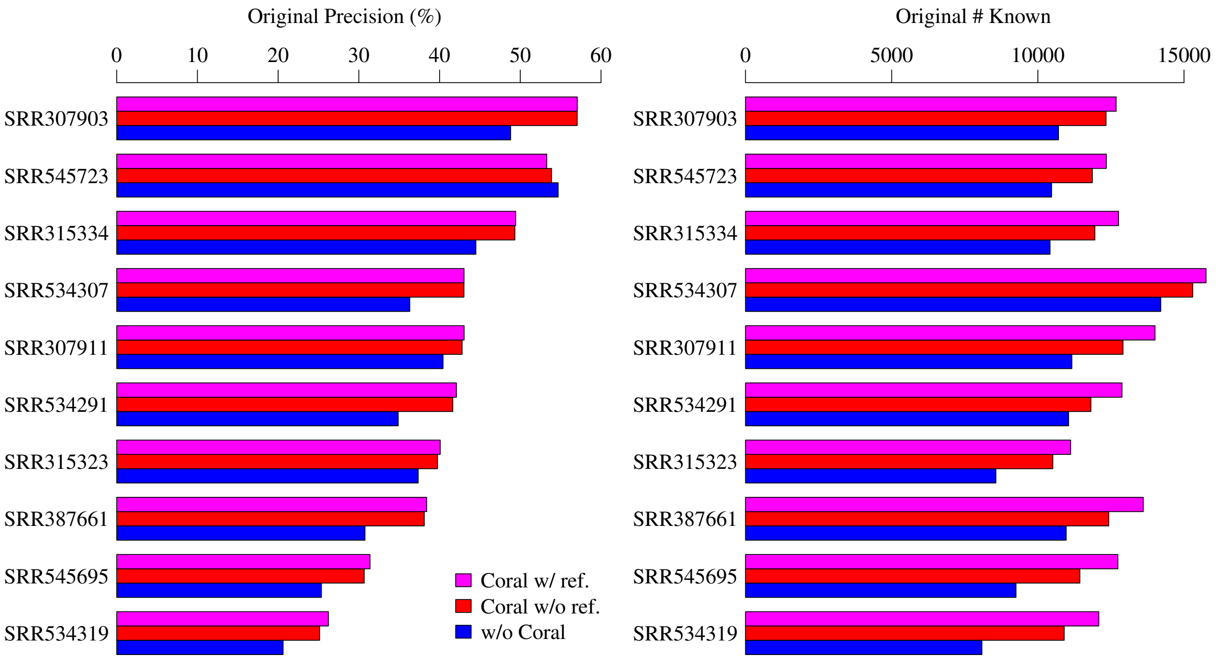


Supplementary Figure 1: Original measures of assembly accuracy comparing the 3 configurations on individual RNA-seq samples of dataset 1. Workflow: Aligner = STAR, Assembler = StringTie.

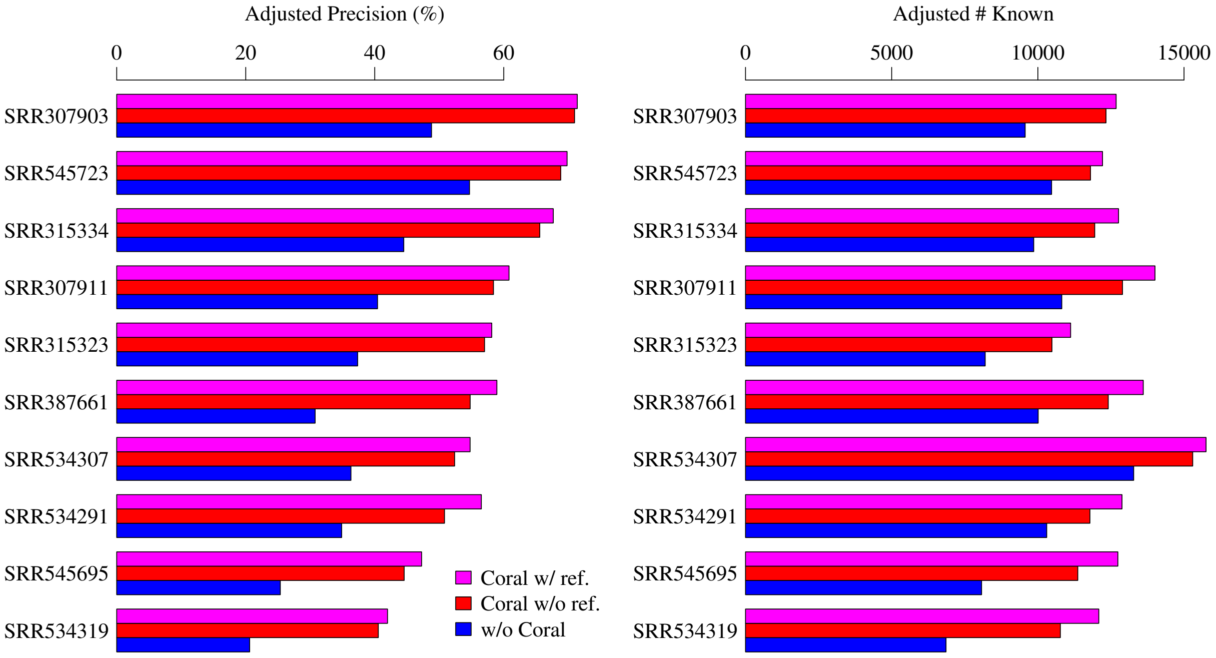


Supplementary Figure 2: Ajuusted measures of assembly accuracy comparing the 3 configurations on individual RNA-seq samples of dataset 1. Workflow: Aligner = STAR, Assembler = StringTie.

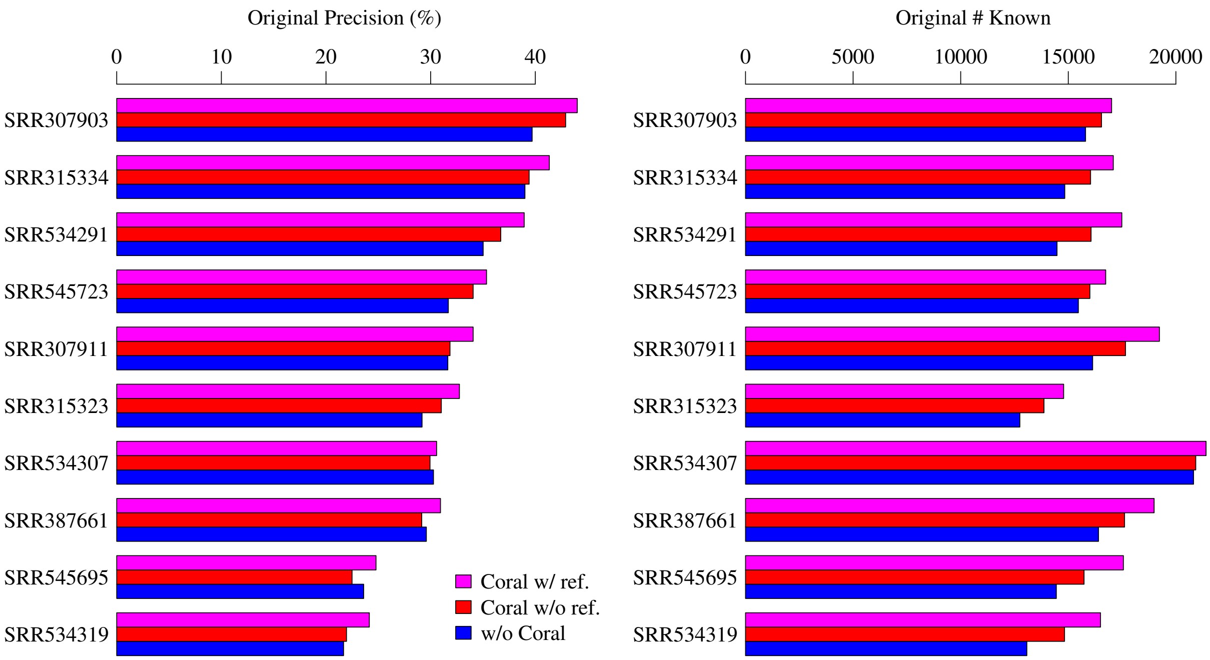


Supplementary Figure 3: Original measures of assembly accuracy comparing the 3 configurations on individual RNA-seq samples of dataset 1. Workflow: Aligner = STAR, Assembler = Scallop.

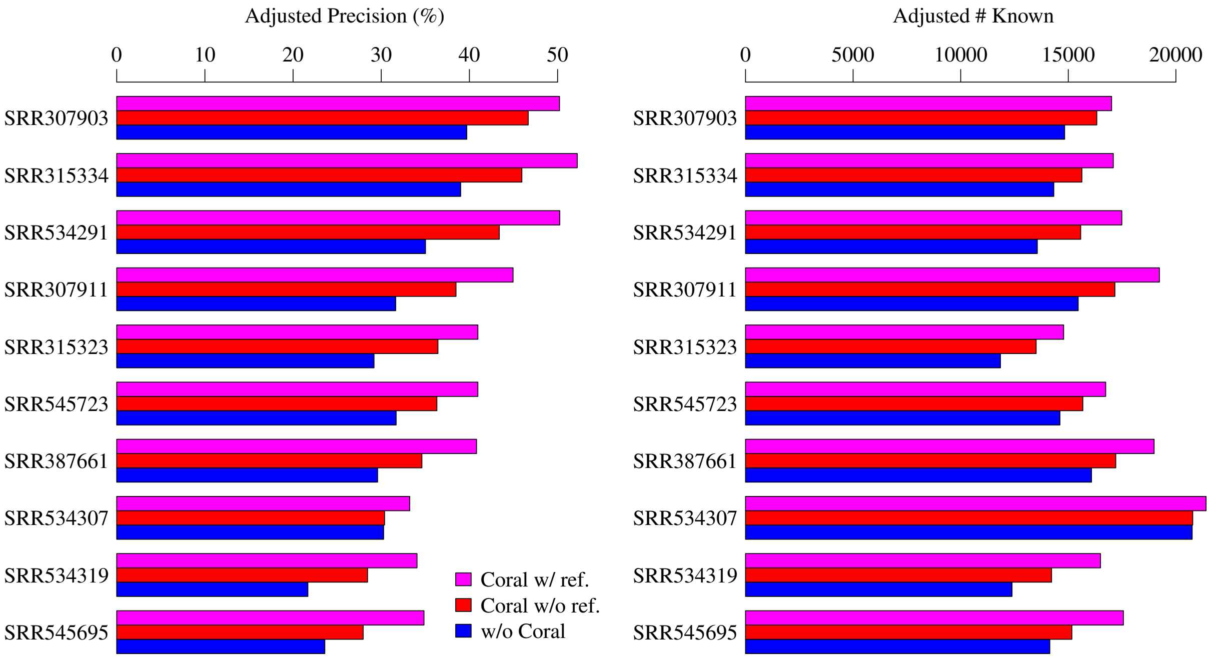


Supplementary Figure 4: Ajuusted measures of assembly accuracy comparing the 3 configurations on individual RNA-seq samples of dataset 1. Workflow: Aligner = STAR, Assembler = Scallop.

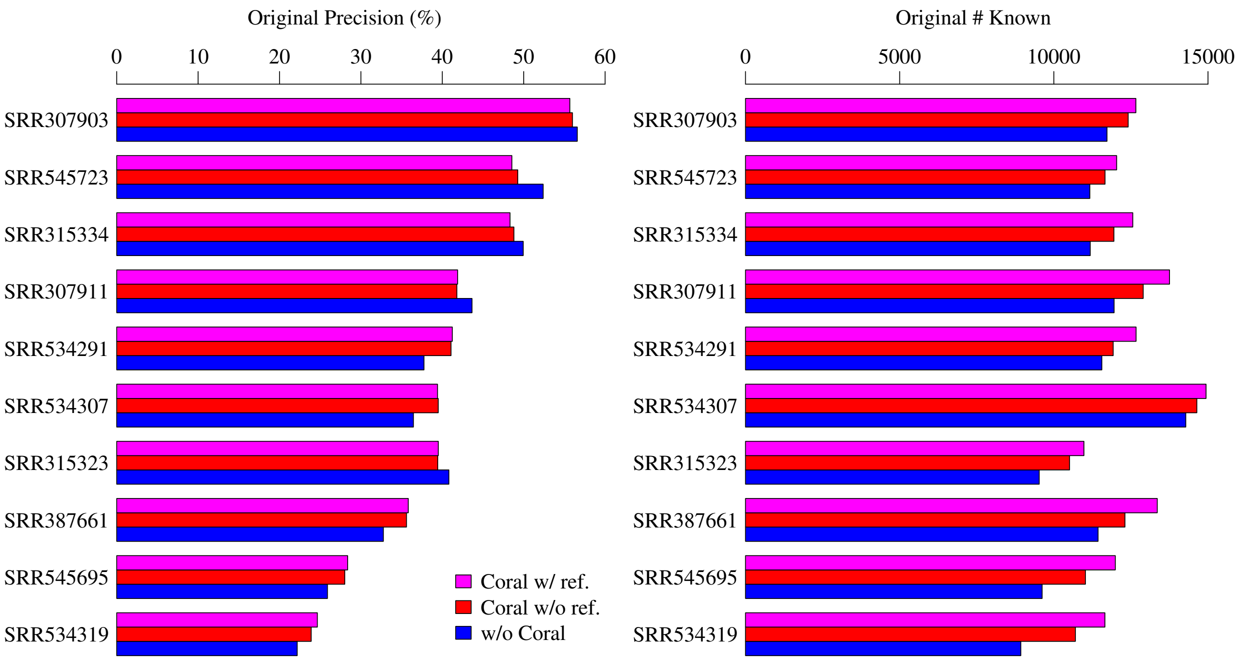


Supplementary Figure 5: Original measures of assembly accuracy comparing the 3 configurations on individual RNA-seq samples of dataset 1. Workflow: Aligner = HISAT, Assembler = StringTie.

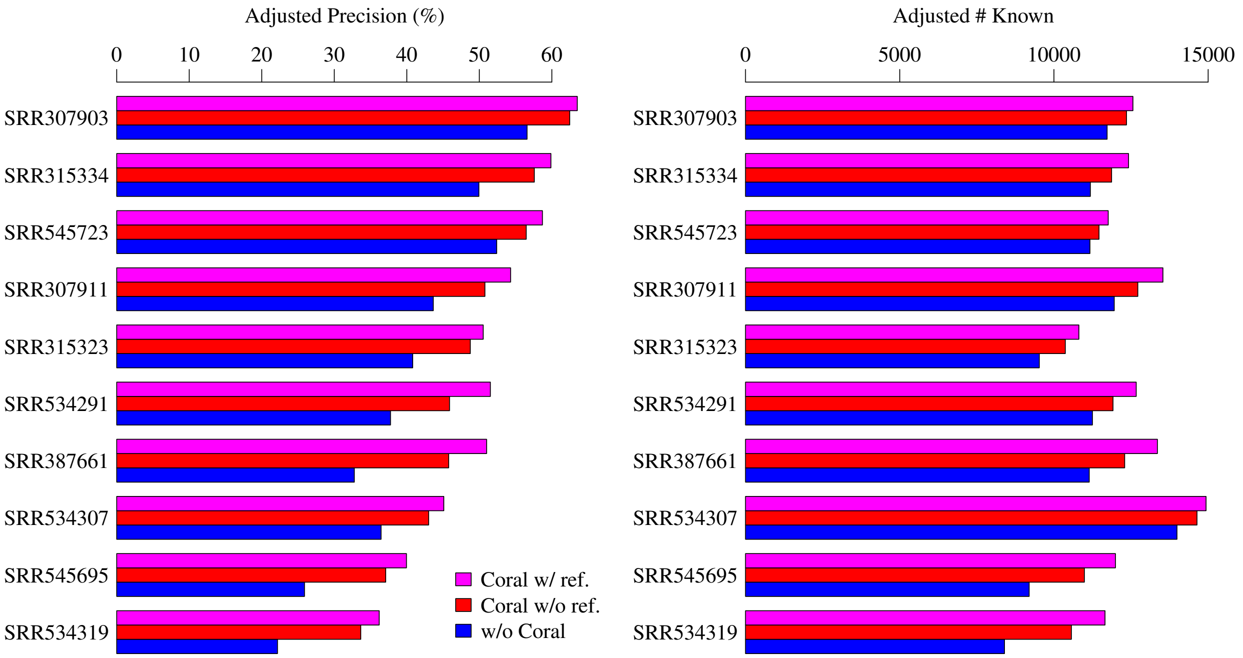


Supplementary Figure 6: Ajuusted measures of assembly accuracy comparing the 3 configurations on individual RNA-seq samples of dataset 1. Workflow: Aligner = HISAT, Assembler = StringTie.

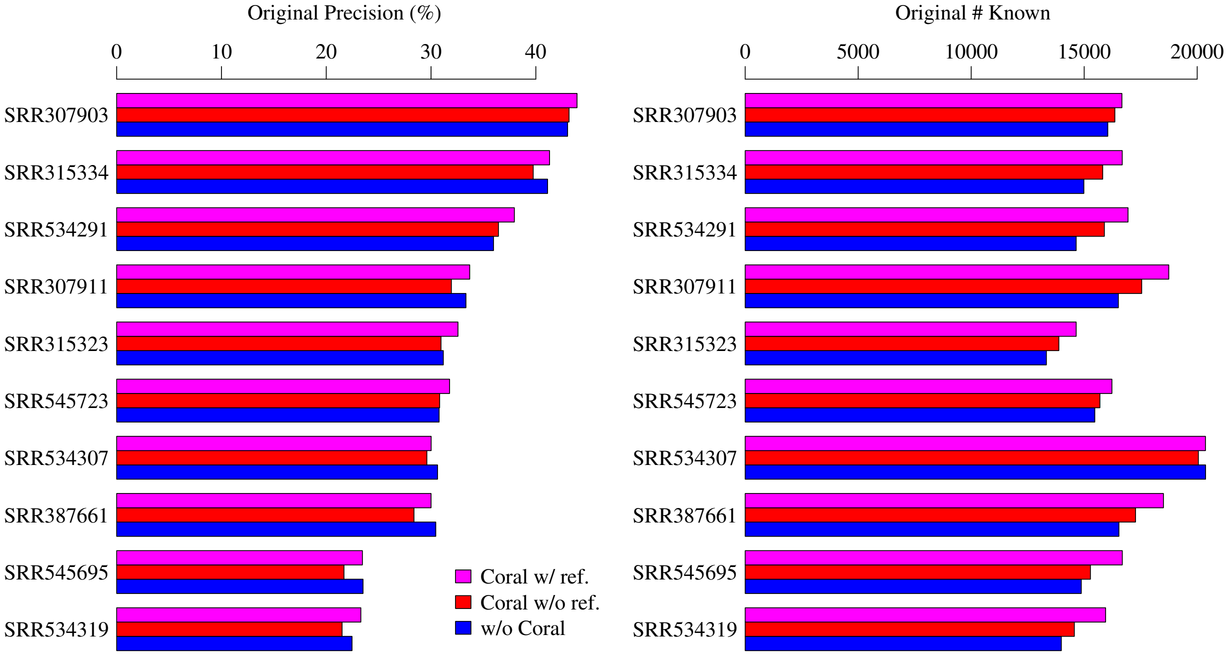


Supplementary Figure 7: Original measures of assembly accuracy comparing the 3 configurations on individual RNA-seq samples of dataset 1. Workflow: Aligner = HISAT, Assembler = Scallop.

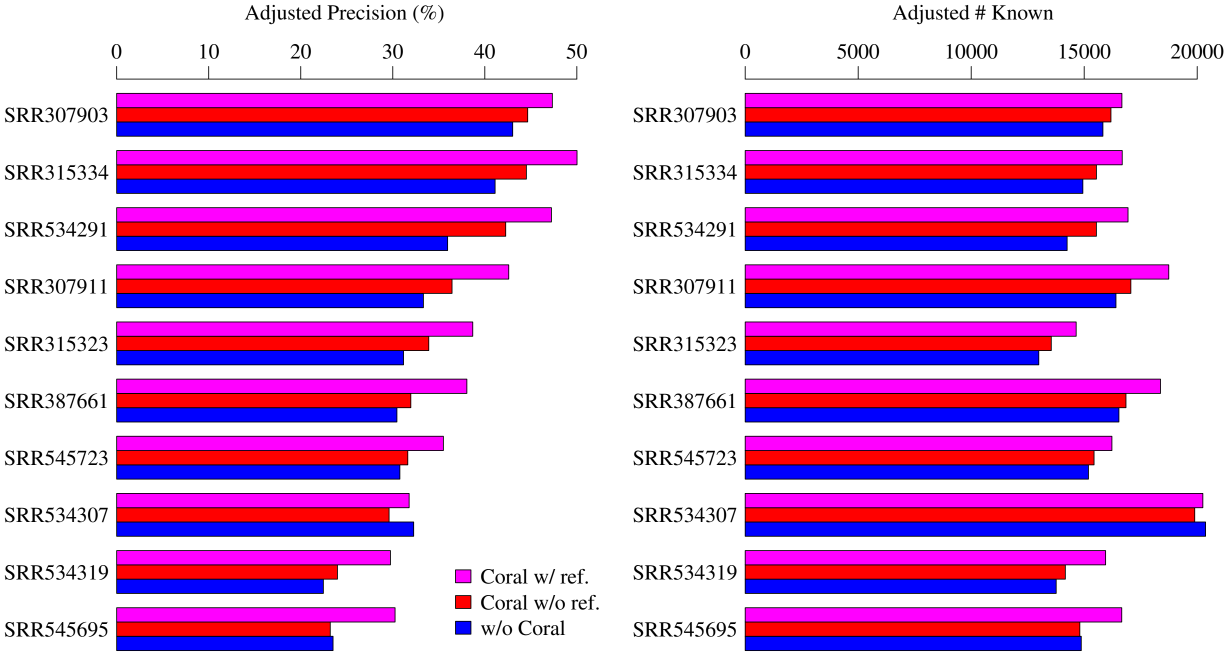


Supplementary Figure 8: Ajuusted measures of assembly accuracy comparing the 3 configurations on individual RNA-seq samples of dataset 1. Workflow: Aligner = HISAT, Assembler = Scallop.

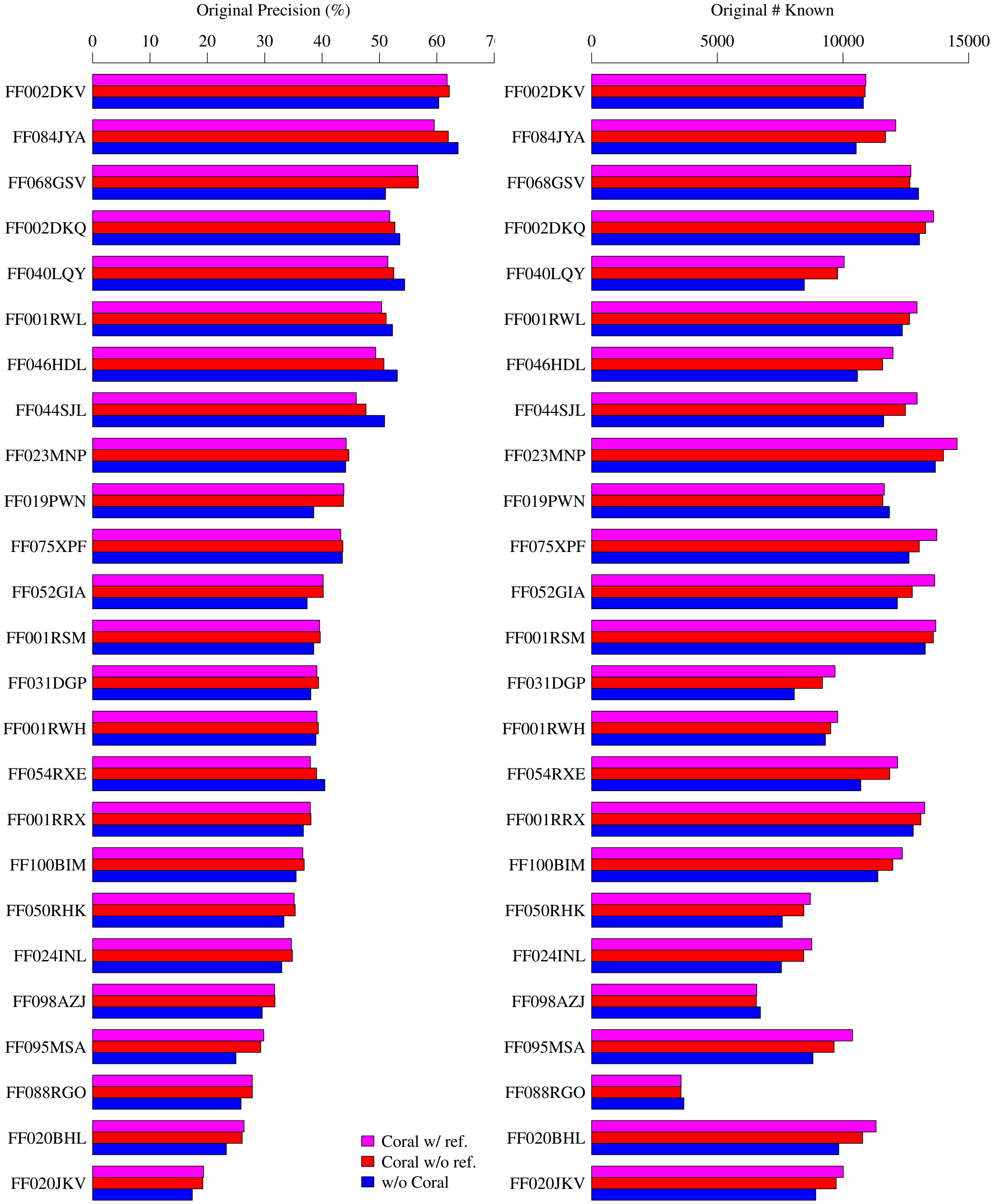


Supplementary Figure 9: Original measures of assembly accuracy comparing the 3 configurations on the first 25 samples of dataset 2. Workflow: Assembler = StringTie.

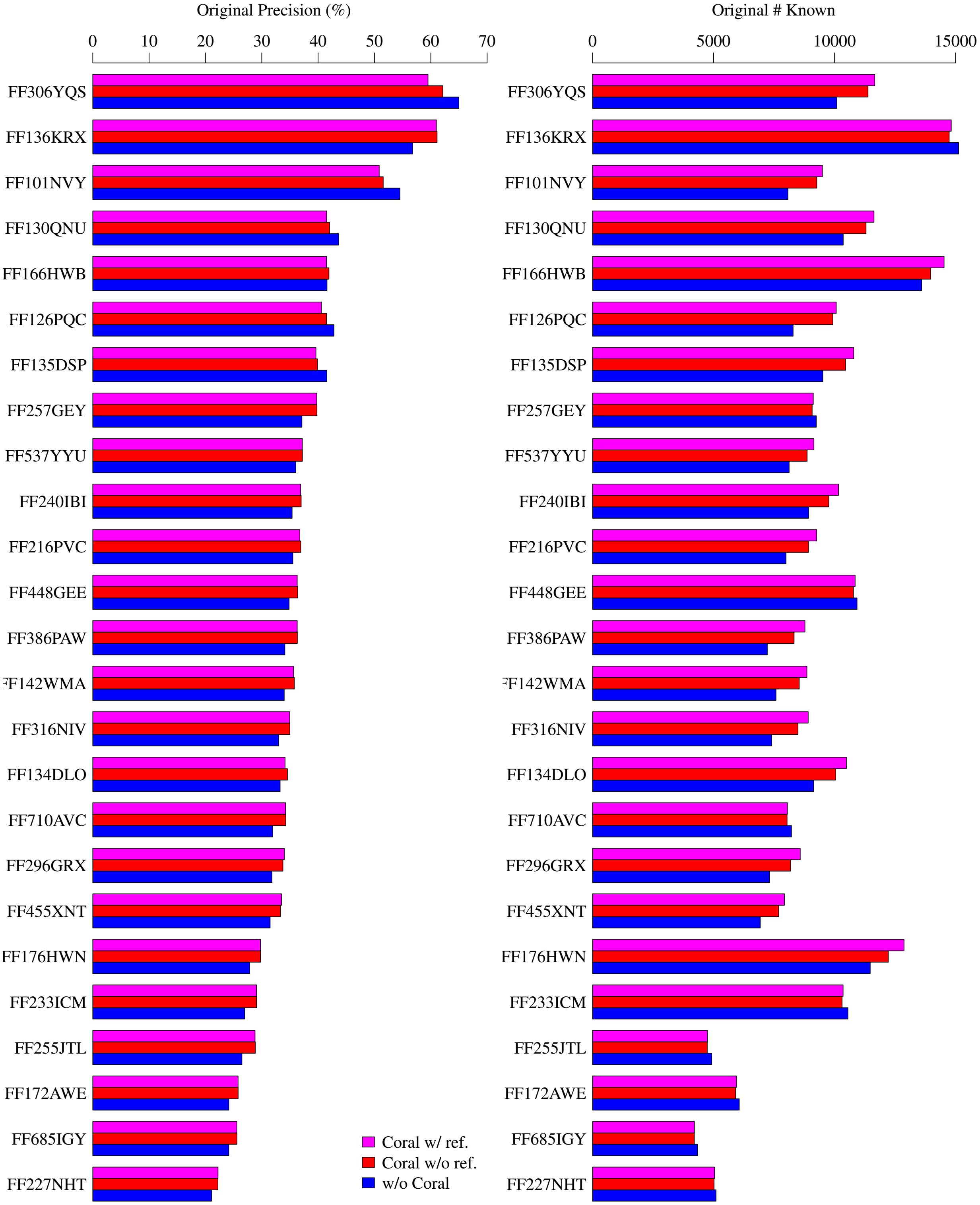


Supplementary Figure 10: Original measures of assembly accuracy comparing the 3 configurations on the second 25 samples of dataset 2. Workflow: Assembler = StringTie.

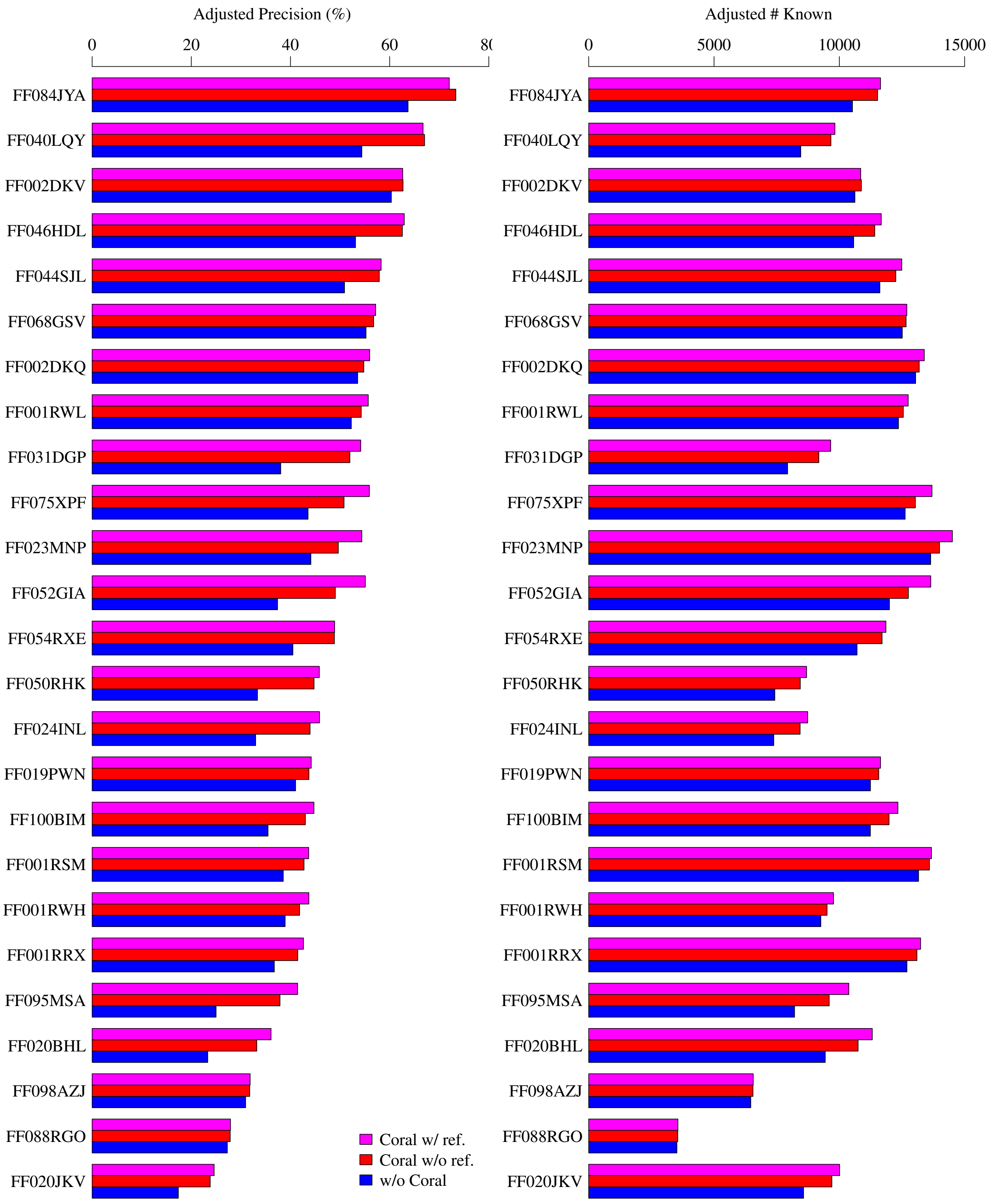


Supplementary Figure 11: Adjusted measures of assembly accuracy comparing the 3 configurations on the first 25 samples of dataset 2. Workflow: Assembler = StringTie.

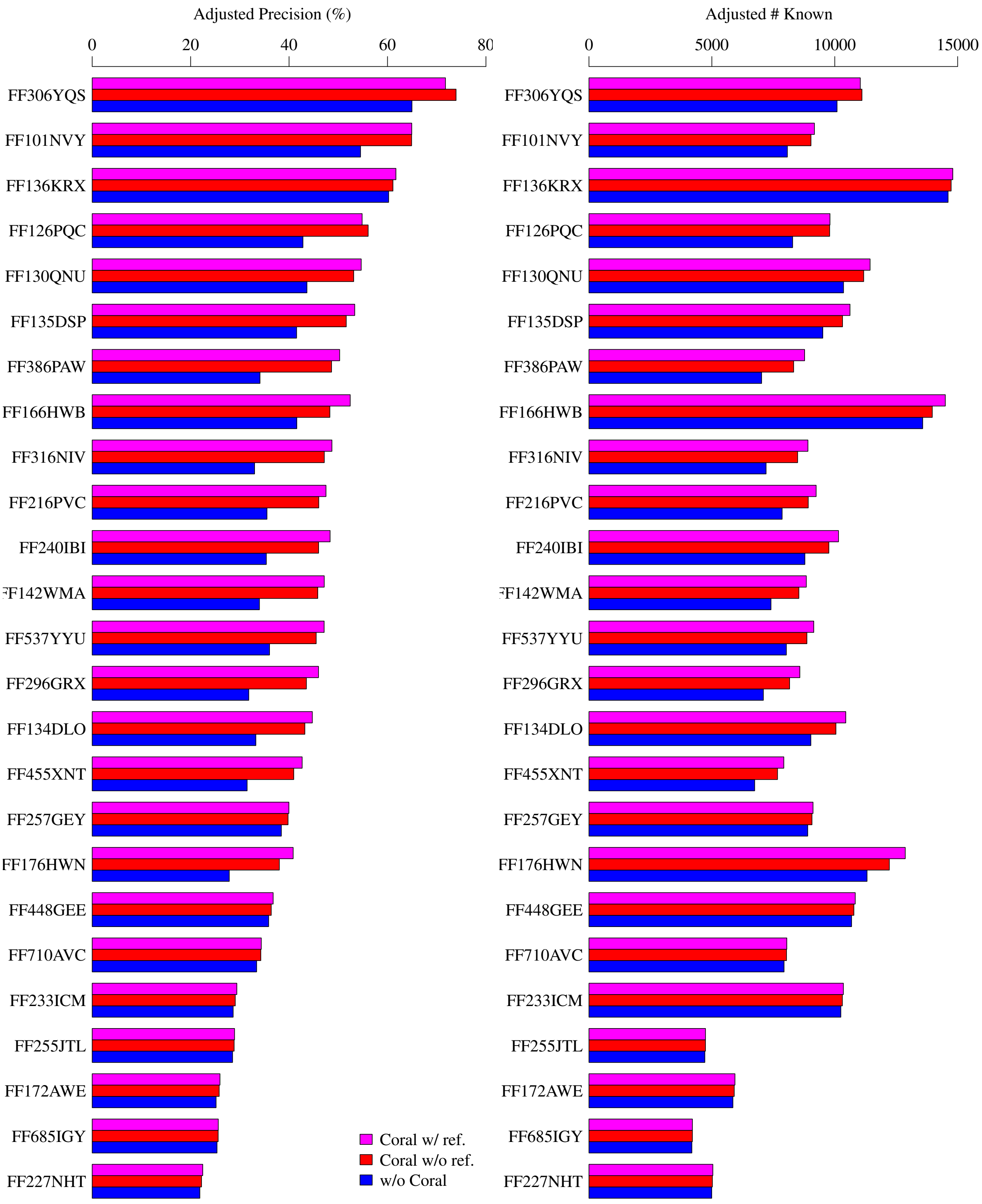


Supplementary Figure 12: Adjusted measures of assembly accuracy comparing the 3 configurations on the second 25 samples of dataset 2. Workflow: Assembler = StringTie.

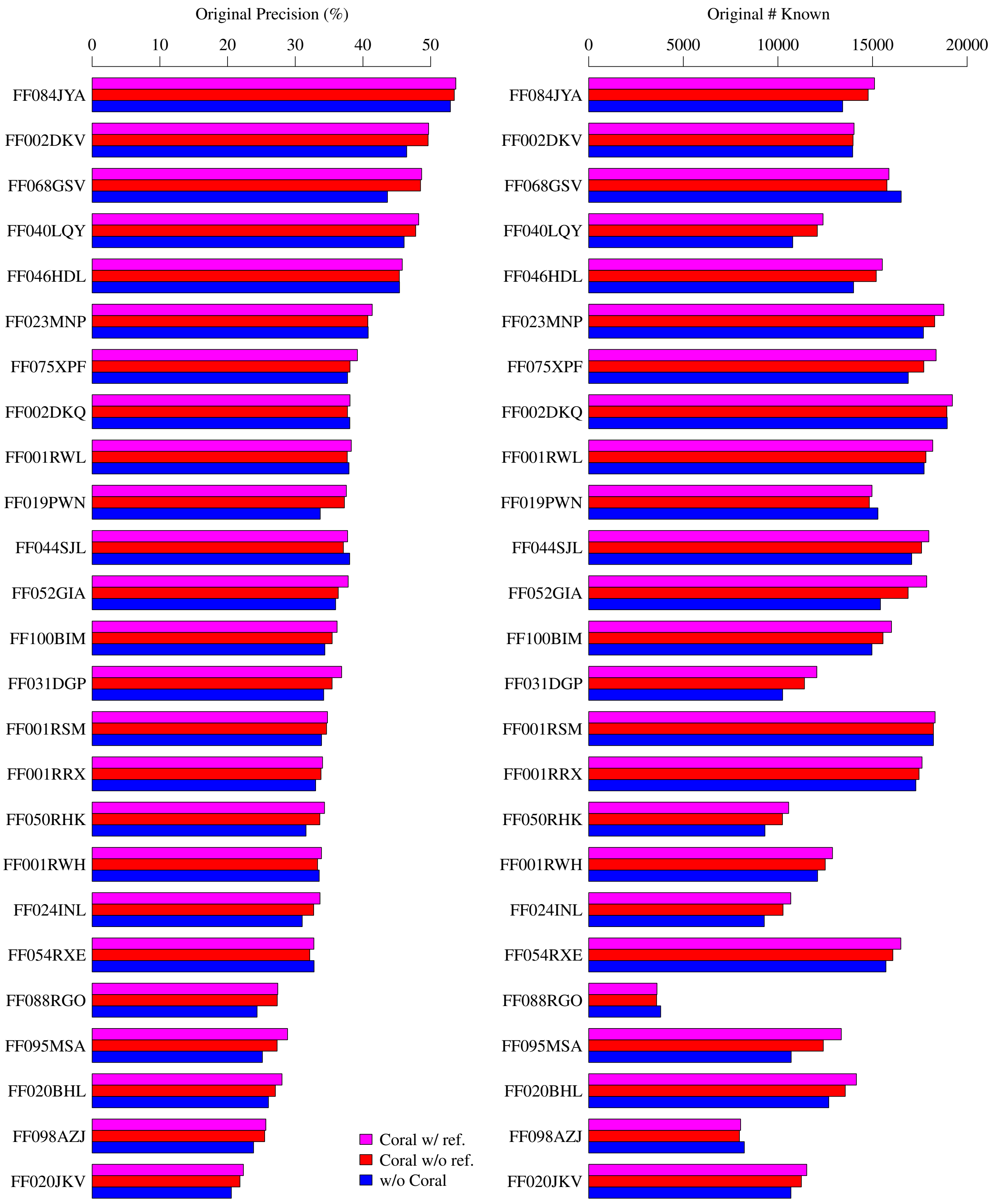


Supplementary Figure 13: Original measures of assembly accuracy comparing the 3 configurations on the first 25 samples of dataset 2. Workflow: Assembler = Scallop.

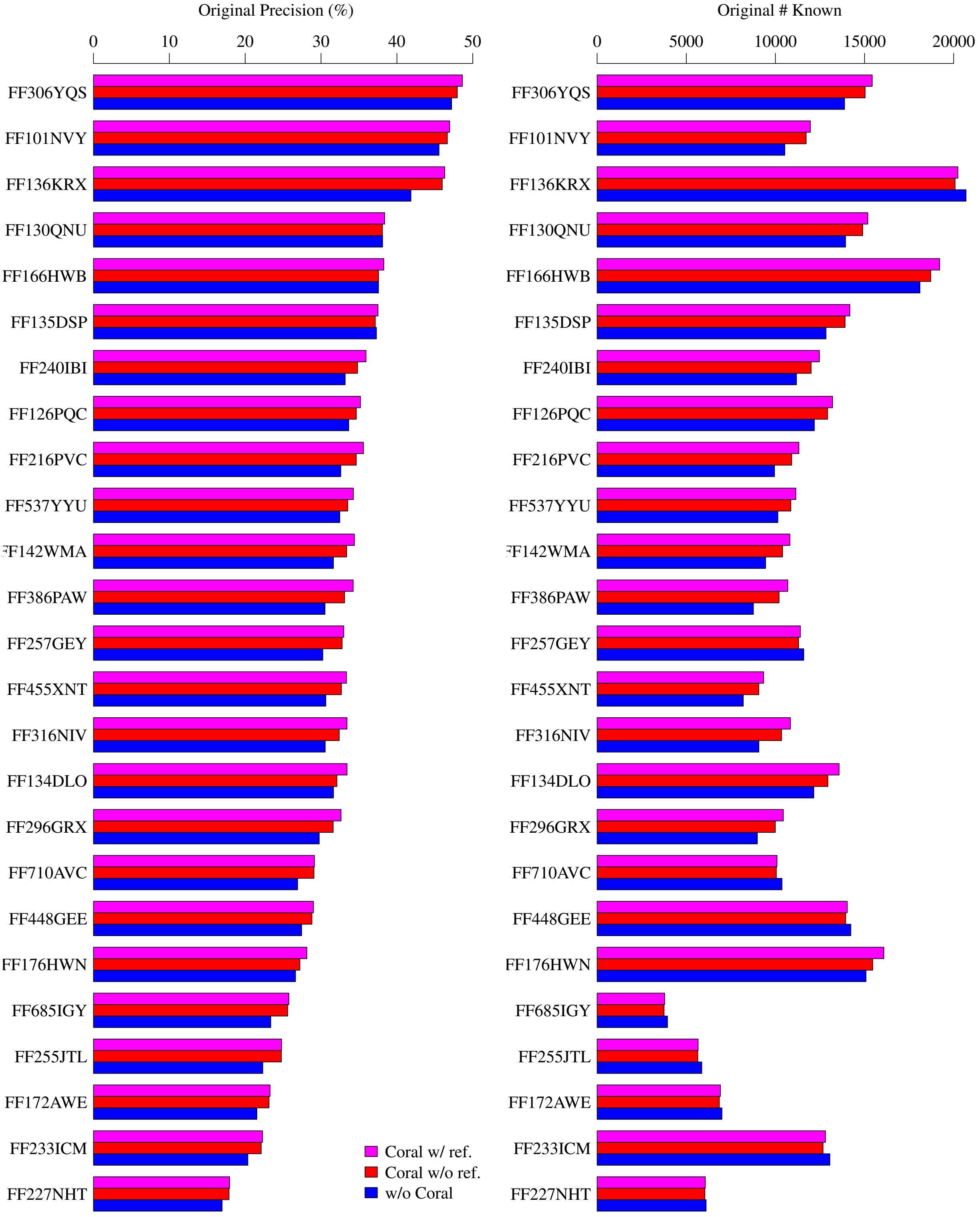


Supplementary Figure 14: Original measures of assembly accuracy comparing the 3 configurations on the second 25 samples of dataset 2. Workflow: Assembler = Scallop.

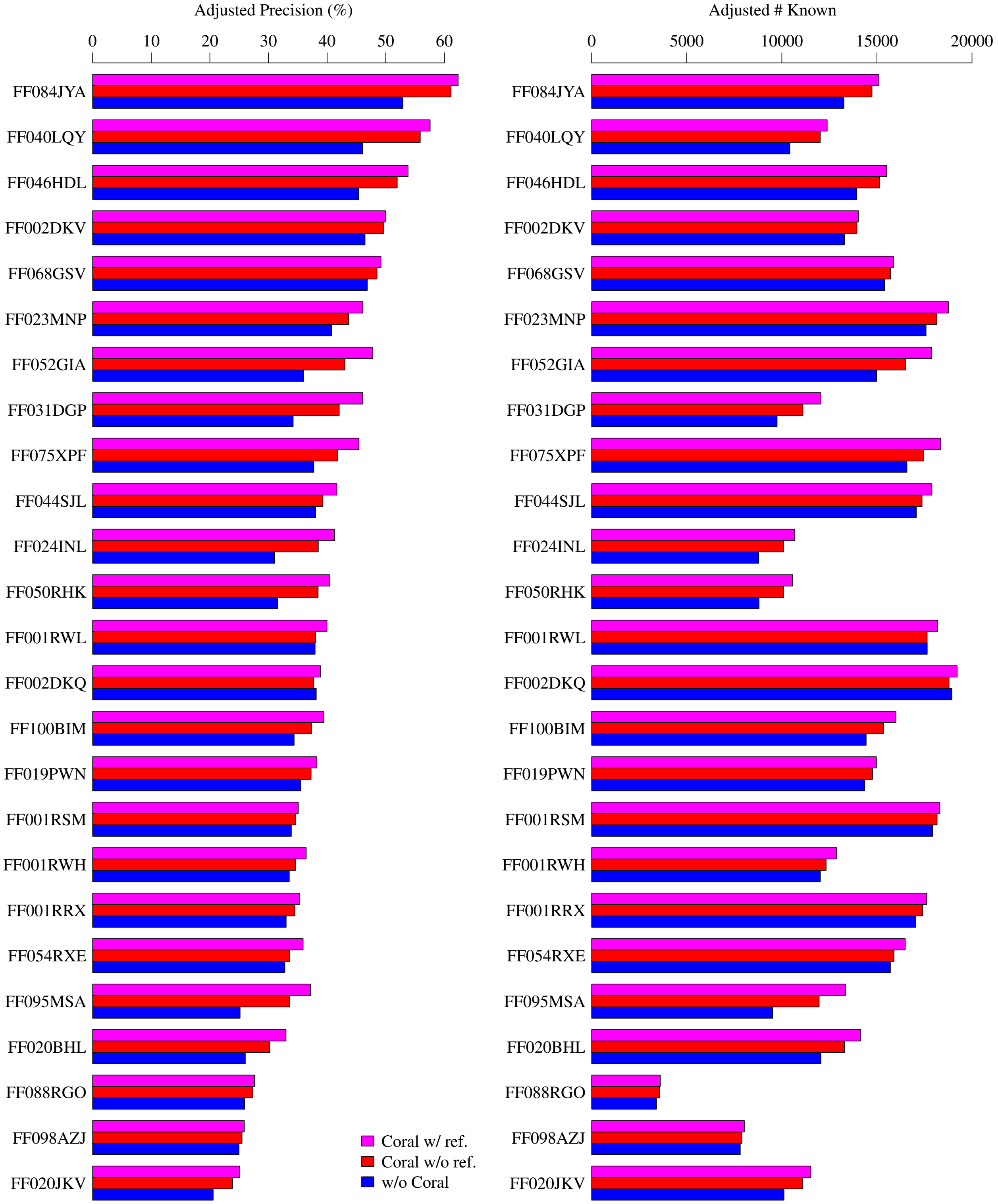


Supplementary Figure 15: Adjusted measures of assembly accuracy comparing the 3 configurations on the first 25 samples of dataset 2. Workflow: Assembler = Scallop.

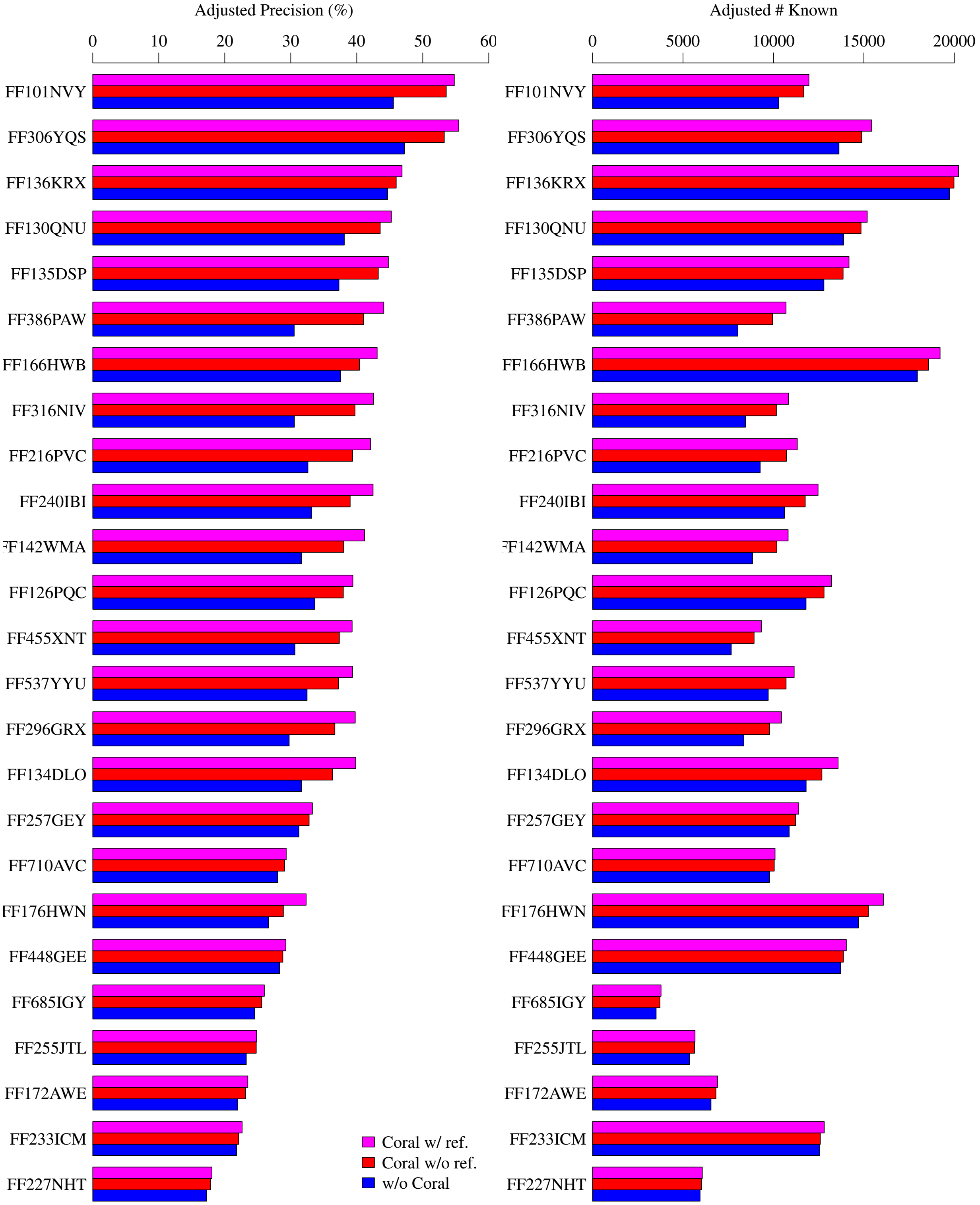


Supplementary Figure 16: Adjusted measures of assembly accuracy comparing the 3 configurations on the second 25 samples of dataset 2. Workflow: Assembler = Scallop.

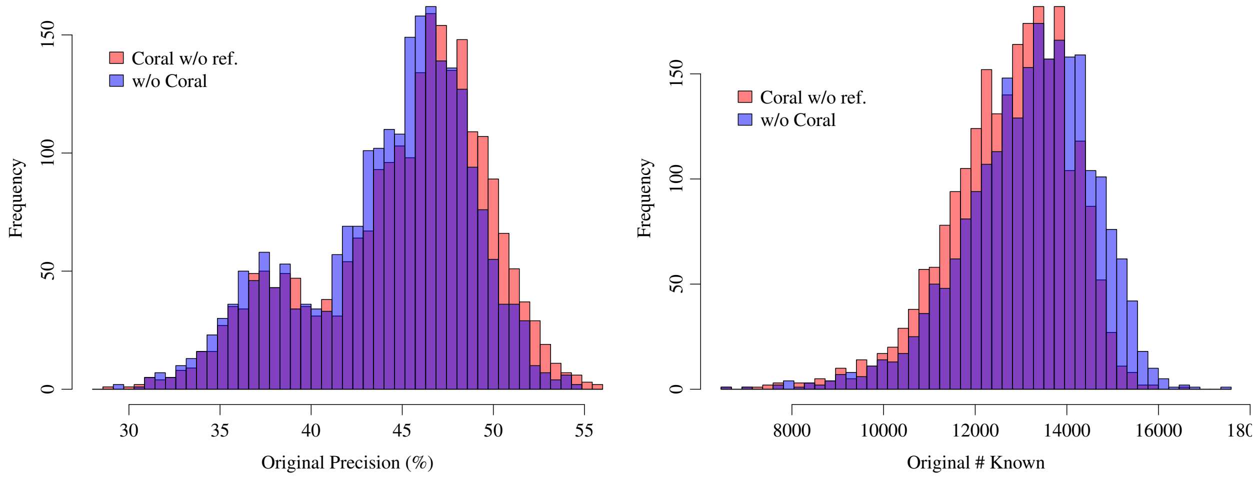


Supplementary Figure 17: Original measures of assembly accuracy comparing the pipeline with and without Coral on samples of dataset 3. Workflow: Assembler = StringTie, and Coral is run without reference annotation (all samples have been pre-aligned in GTEx and we directly use them).

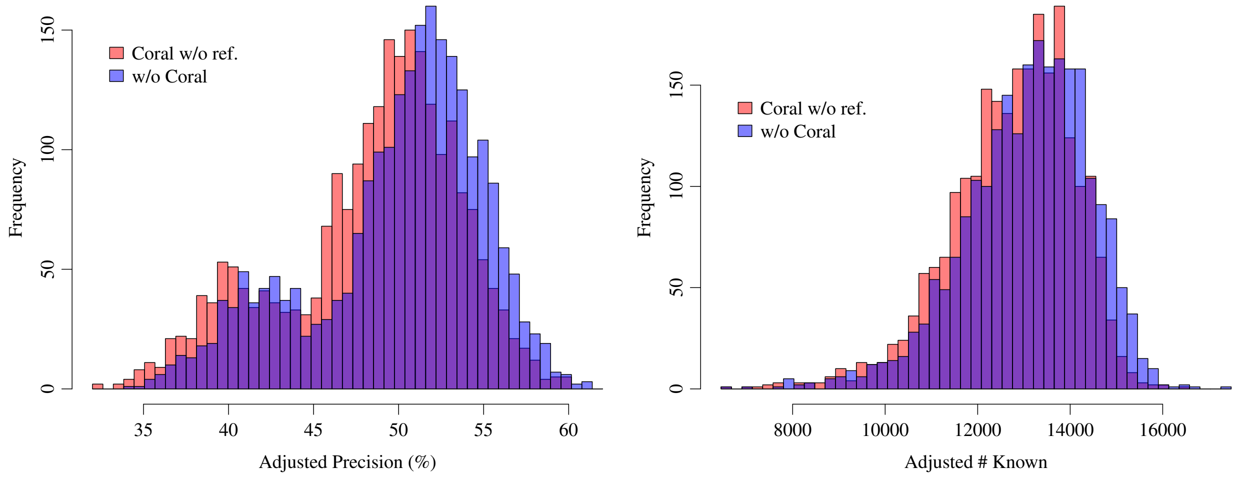


Supplementary Figure 18: Adjusted measures of assembly accuracy comparing the pipeline with and without Coral on samples of dataset 3. Workflow: Assembler = StringTie, and Coral is run without reference annotation (all samples have been pre-aligned in GTEx and we directly use them).

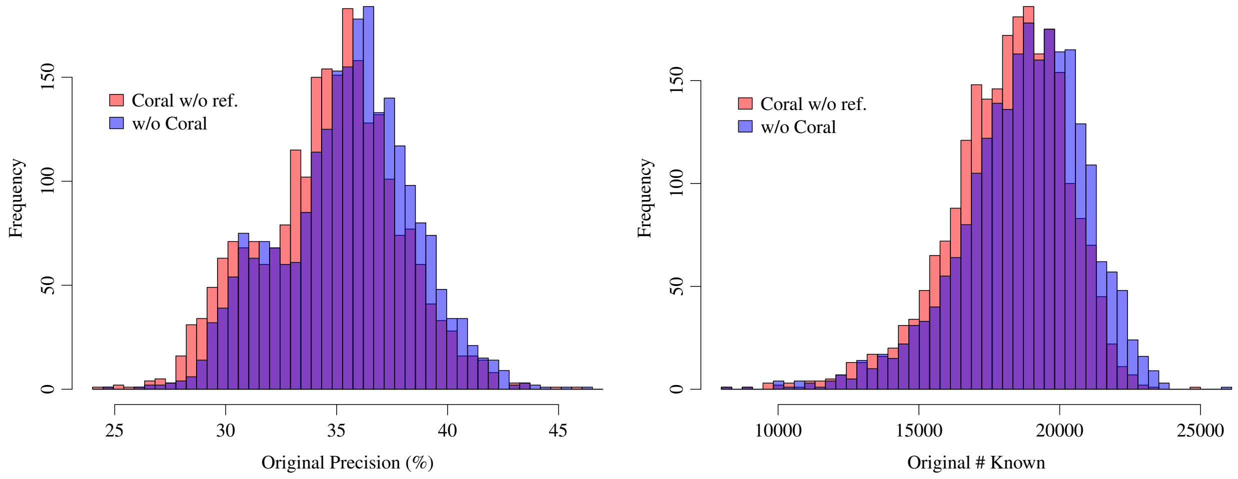


Supplementary Figure 19: Original measures of assembly accuracy comparing the pipeline with and without Coral on samples of dataset 3. Workflow: Assembler = Scallop, and Coral is run without reference annotation (all samples have been pre-aligned in GTEx and we directly use them).

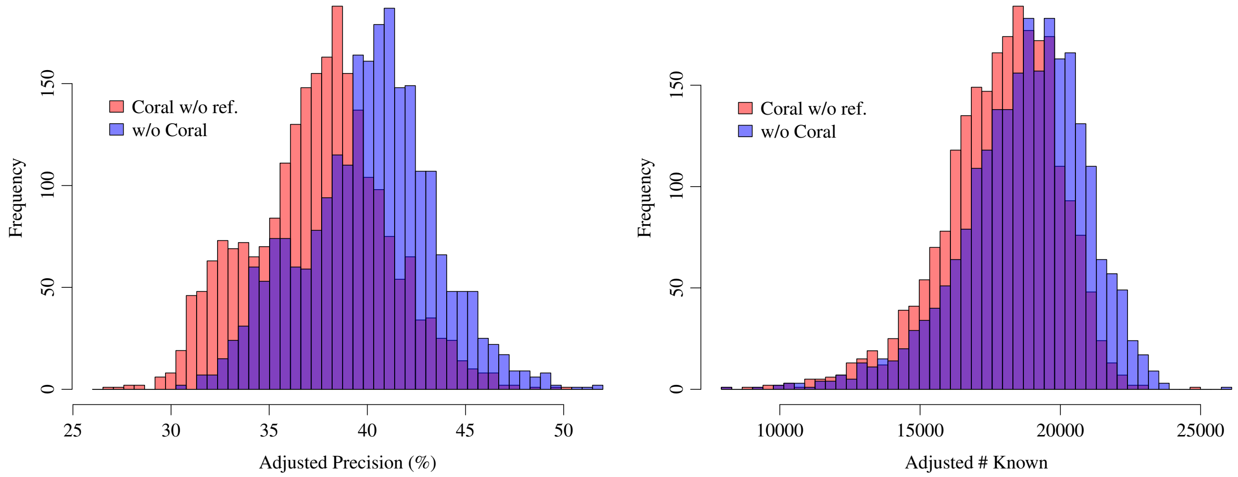


Supplementary Figure 20: Adjusted measures of assembly accuracy comparing the pipeline with and without Coral on samples of dataset 3. Workflow: Assembler = Scallop, and Coral is run without reference annotation (all samples have been pre-aligned in GTEx and we directly use them).

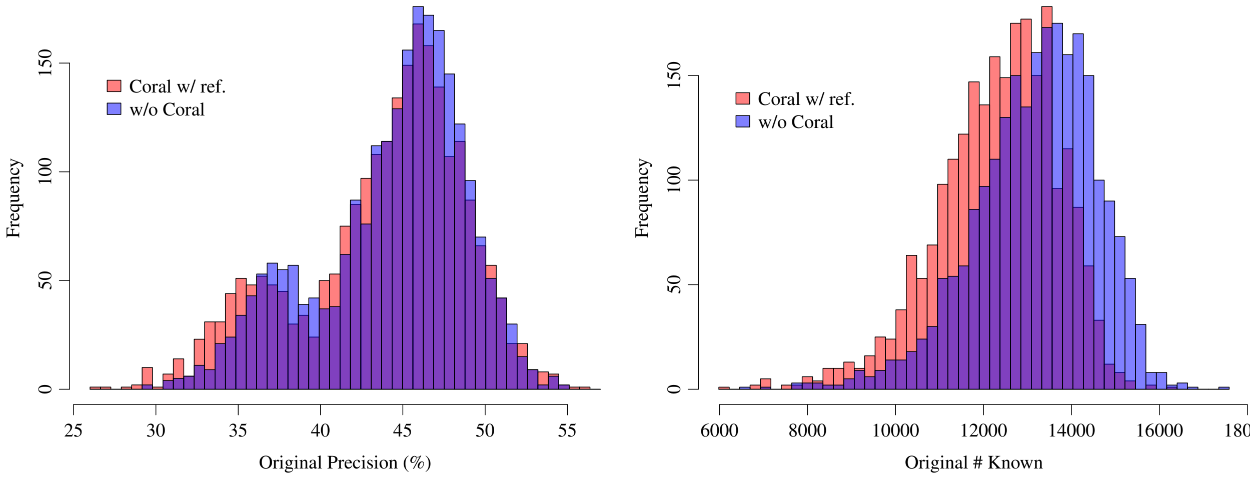


Supplementary Figure 21: Original measures of assembly accuracy comparing the pipeline with and without Coral on samples of dataset 3. Workflow: Assembler = StringTie, and Coral is run without reference annotation (all samples have been pre-aligned in GTEx and we directly use them).

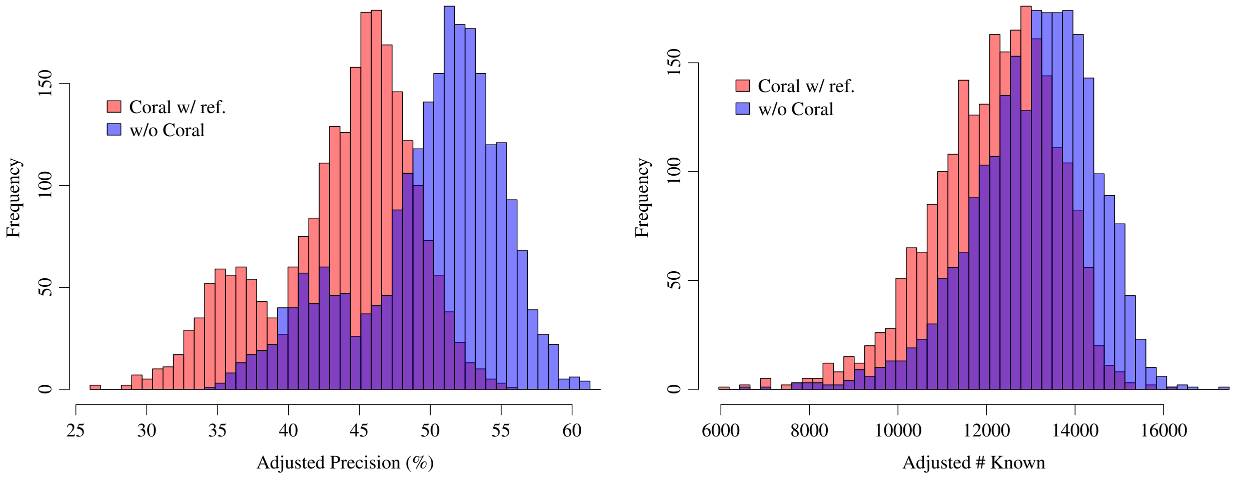


Supplementary Figure 22: Adjusted measures of assembly accuracy comparing the pipeline with and without Coral on samples of dataset 3. Workflow: Assembler = StringTie, and Coral is run without reference annotation (all samples have been pre-aligned in GTEx and we directly use them).

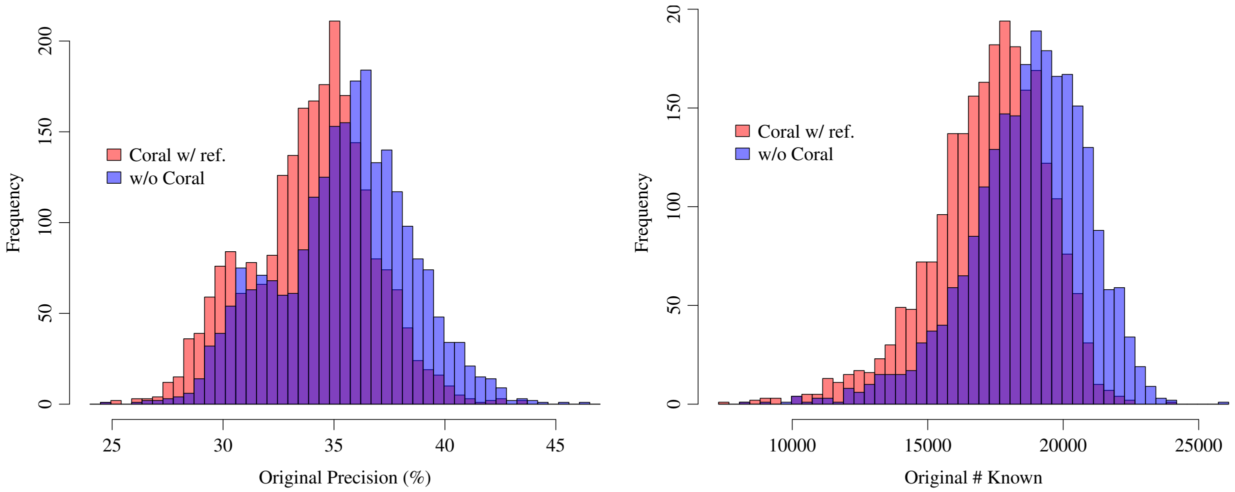


Supplementary Figure 23: Original measures of assembly accuracy comparing the pipeline with and without Coral on samples of dataset 3. Workflow: Assembler = Scallop, and Coral is run without reference annotation (all samples have been pre-aligned in GTEx and we directly use them).

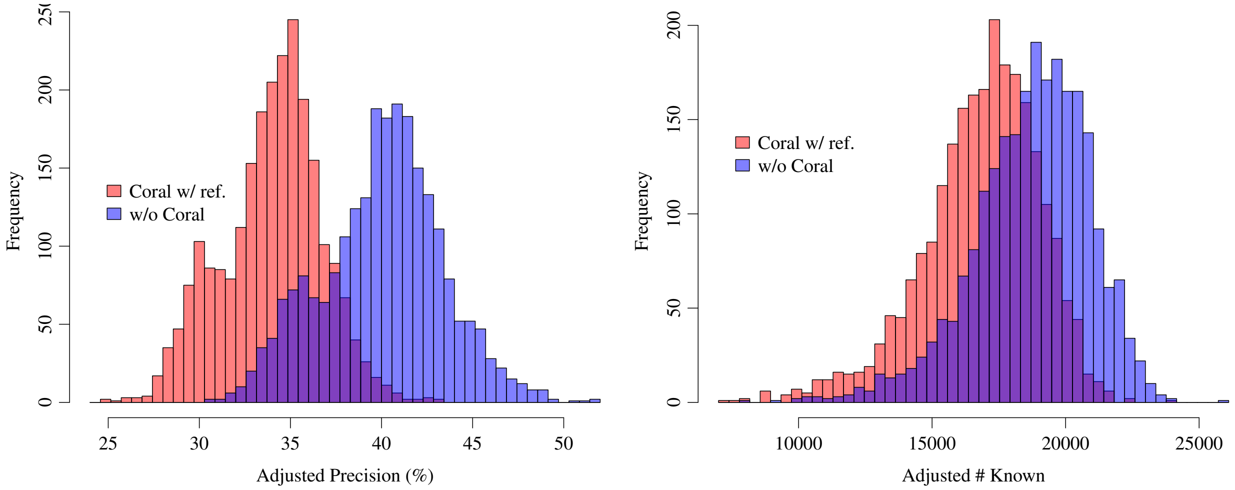


Supplementary Figure 24: Adjusted measures of assembly accuracy comparing the pipeline with and without Coral on samples of dataset 3. Workflow: Assembler = Scallop, and Coral is run without reference annotation (all samples have been pre-aligned in GTEx and we directly use them).
